# Profile-Wise Analysis: A profile likelihood-based workflow for identifiability analysis, estimation, and prediction with mechanistic mathematical models

**DOI:** 10.1101/2022.12.14.520367

**Authors:** Matthew J. Simpson, Oliver J. Maclaren

**Affiliations:** School of Mathematical Sciences, Queensland University of Technology, Brisbane, Queensland 4001, Australia; Department of Engineering Science, University of Auckland, Auckland 1142, New Zealand

## Abstract

Interpreting data using mechanistic mathematical models provides a foundation for discovery and decision-making in all areas of science and engineering. Developing mechanistic insight by combining mathematical models and experimental data is especially critical in mathematical biology as new data and new types of data are collected and reported. Key steps in using mechanistic mathematical models to interpret data include: (i) identifiability analysis; (ii) parameter estimation; and (iii) model prediction. Here we present a systematic, computationally-efficient workflow we call *Profile-Wise Analysis* (PWA) that addresses all three steps in a unified way. Recently-developed methods for constructing ‘profile-wise’ prediction intervals enable this workflow and provide the central linkage between different workflow components. These methods propagate profile-likelihood-based confidence sets for model parameters to predictions in a way that isolates how different parameter combinations affect model predictions. We show how to extend these profile-wise prediction intervals to two-dimensional interest parameters. We then demonstrate how to combine profile-wise prediction confidence sets to give an overall prediction confidence set that approximates the full likelihood-based prediction confidence set well. Our three case studies illustrate practical aspects of the workflow, focusing on ordinary differential equation (ODE) mechanistic models with both Gaussian and non-Gaussian noise models. While the case studies focus on ODE-based models, the workflow applies to other classes of mathematical models, including partial differential equations and simulation-based stochastic models. Open-source software on GitHub can be used to replicate the case studies.

## 1 Introduction

Mechanistic mathematical models, typically represented using differential equations or other evolution equations, underpin scientific understanding and thus play an essential role in science and engineering. However, developing and solving a mathematical or computational model is only the first step in realizing its potential value: we must reliably connect mathematical models to empirical data to ensure their relevance. There have been significant recent efforts to develop ‘Bayesian workflows’ [1–7] for systematizing these tasks in a Bayesian setting, but much less attention on corresponding frequentist workflows, and particularly so for frequentist approaches incorporating prediction. Although both Bayesian and frequentist approaches are concerned with uncertainty quantification in the general sense, the goals, interpretation and computational requirements of frequentist and Bayesian approaches to this task are different [8, 9]. In terms of goals and interpretation, frequentist methods are generally concerned with constructing estimation procedures with reliability guarantees (such as ‘coverage’ of a confidence interval) under repeated use in ‘naturally random’ environments modelled by probability distributions [8–10]. In contrast, Bayesian methods generally aim to ensure the ‘coherence’ of an analyst’s personal beliefs or ‘state of information’ when updating them given new data, representing the analyst’s beliefs (rather than the external environment) by a probability distribution [8, 9, 11, 12]. Computationally, Bayesian methods are typically more expensive; as one example, see, e.g. [13]. However, this assessment depends on the computational techniques used (e.g. sampling or optimization-based) and end targets of the analysis (e.g. point estimates, set-valued estimates, or distribution-valued estimates).

Here we aim to provide an efficient, likelihood-based frequentist workflow for systematic and efficient use of data to understand and predict with mechanistic mathematical models. The new workflow, called *Profile-Wise Analysis* (PWA), enables parameter identifiability analysis, mechanistic-model-based parameter estimation, analysis of the sensitivity of predictions with respect to different parameters and different parameter combinations, as well as the construction of confidence intervals for predictive quantities. Our current approach is explicitly frequentist in motivation; however, we take no position on whether frequentist or Bayesian inference is inherently more desirable than the other, only that they have different goals and interpretations, are appropriate for their respective goals, and can thus provide complementary rather than directly competing analyses [8].

### Existing approaches for uncertainty propagation and prediction

As discussed in the next section, ‘New approach: Profile-Wise Analysis’, frequentist tools for identifiability analysis and parameter estimation are well-established in systems biology and related fields. In contrast, tools for frequentist prediction uncertainty for mechanistic models are less well-developed; hence, we focus on this literature in the present section. As with parameter estimation [8], the logic of propagation of uncertainty from parameters to predictions is different in frequentist inference and Bayesian inference (see e.g. [14]). Hence Bayesian uncertainty propagation methods are not directly applicable to frequentist inference, and we require approaches tailored to frequentist goals. We describe the logic of uncertainty propagation in a frequentist setting more explicitly when describing our methodology later; for now, we review existing methods.

The current state of frequentist prediction for mechanistic models is well-summarised by Villaverde et al. [15] who write, in a recent review in the context of systems biology models:

> Unfortunately, estimating prediction uncertainties accurately is nontrivial, due to the nonlinear dependence of model characteristics on parameters. While a number of numerical approaches have been proposed for this task, their strengths and weaknesses have not been systematically assessed yet.

Villaverde et al. then compare different predictive uncertainty quantification methods in systems biology. These include the frequentist methods of local linearization and covariance (i.e. Fisher information-based) propagation [16], prediction profile likelihood [17–19], and ensemble methods [20]. We consider this latter set of methods to include bootstrap-style methods such as the bootstrap version of randomized maximum likelihood (RML) [21] we presented recently alongside profile-based methods in [22].

Villaverde et al. [15] conclude that existing methods involve trade-offs between computational tractability and statistical rigor. While the so-called profile prediction likelihood [19] is, in principle, justified rigorously and directly by standard profile likelihood theory [18, 19, 22–24], this approach is challenging to implement and computationally expensive. Linearization is simple and efficient but potentially imprecise for highly-nonlinear models. Ensemble methods are simple to implement but also typically expensive and, depending on the specific form, can be ad hoc and challenging to analyze in terms of accuracy [15].

As in the profile prediction likelihood approach [17–19], our workflow, including the prediction component, is based on profile likelihood theory. However, our approach is much simpler to implement and interpret as, in contrast to existing methods [17–19], we use profile-likelihood for prediction in a new way that avoids specifying additional optimal control or integration-based steps. We intentionally compare our method to a rigorously justifiable but inefficient method: direct Monte Carlo evaluation and propagation (post-thresholding) of the full likelihood. Our workflow thus provides a new frequentist methodology for prediction using mechanistic models, alongside identifiability analysis and estimation. We evaluate the new workflow against a ‘gold-standard’ likelihood-based approach, which is rigorous but inefficient (we explicitly justify the classification of this approach as gold-standard in the Results section). In the interests of fair comparison and exposition, we focus on canonical mechanistic models here. However, in addition to providing a modular component of a systematic workflow for understanding the influence of various parameters and parameter combinations, our prediction method is conceptually and computationally more straightforward and less expensive than existing profile-likelihood-based prediction methods and is designed with future scalability in mind. Furthermore, as our approach retains the link between finite-dimensional model parameter vectors and ‘infinite-dimensional’ model trajectories, it naturally provides fully ‘curvewise’ (simultaneous) predictive confidence sets for model trajectories, in contrast to the typical ‘pointwise’ intervals of other methods [17–19]. This means we expect our 95% confidence sets for the trajectory to trap the full trajectory 95% of the time, which is a stronger and often more relevant guarantee than merely trapping the trajectory at particular time points separately [64].

### New approach: Profile-Wise Analysis

Here we provide a brief overview of our profile likelihood-based workflow, *Profile-Wise Analysis* (PWA), its key components, and relation to the literature. Figure 1 summarises this workflow. The Results section contains the full details and definitions whereas here we outline some background information.

**Figure 1:**
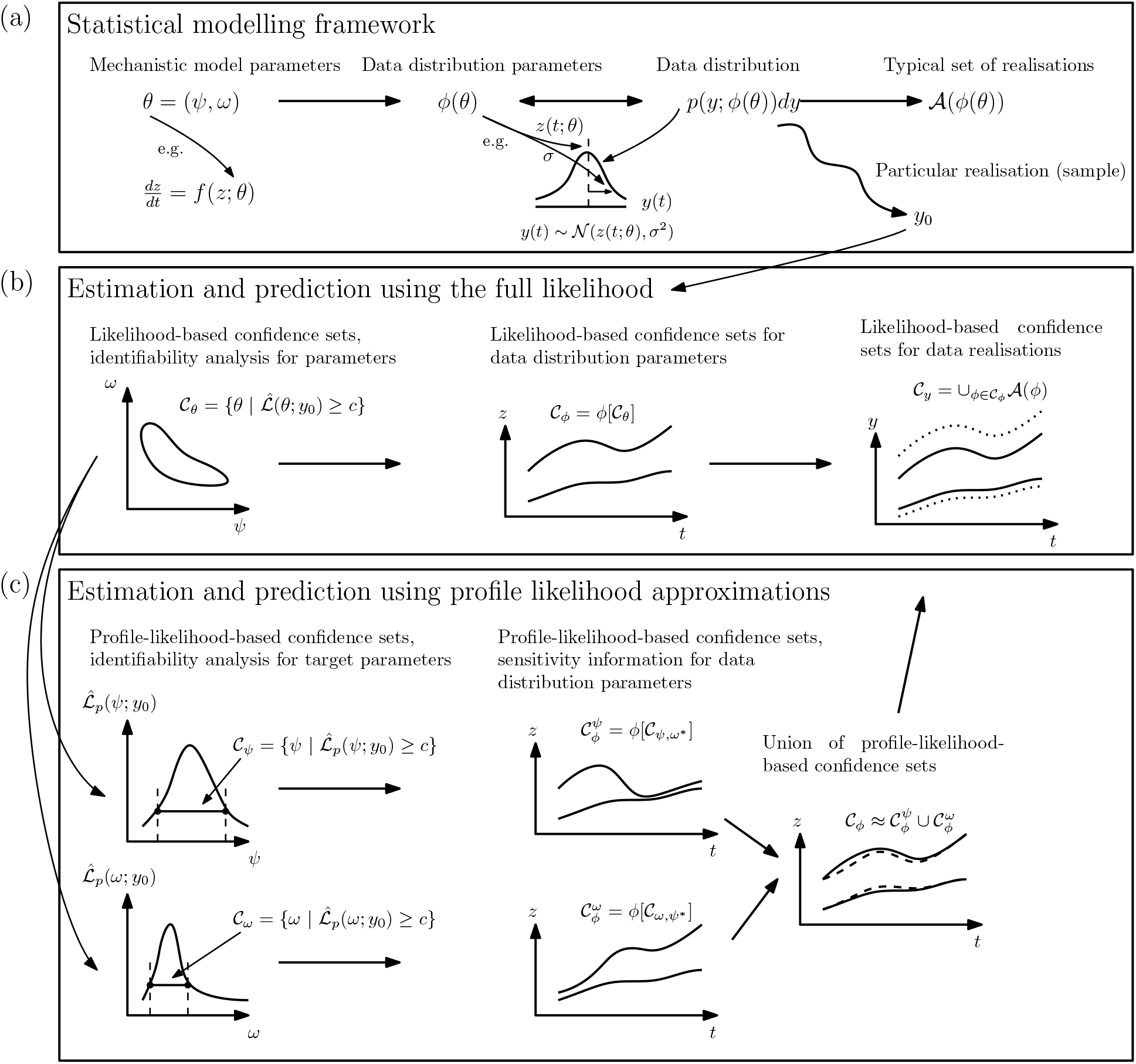
Profile-Wise Analysis: A profile likelihood-based workflow illustrating: (a) statistical framework; (b) estimation and prediction using the full likelihood function; and, (c) estimation and prediction using profile-wise, profile likelihood approximations.

#### Likelihood functions and auxiliary mappings

As our workflow uses an explicitly likelihood-based approach to statistical inference, we need to have a likelihood function available. As detailed in the Results section, a generic likelihood function is based on a probability model of the data given the parameters of the form

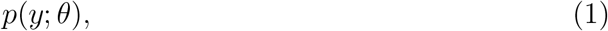

for mechanistic model parameters *θ* and data *y*. This probability model can be viewed as a mapping, *θ 1→ p*(*y*; *θ*), corresponding to the so-called ‘forward mapping’ in the field of inverse problems [25, 26]. The likelihood function is then obtained from this model by treating this as a function of the parameter for fixed data [9, 27, 28] (see Results).

Having an explicit form for the forward mapping, and hence for the likelihood, is typical for statistical models with the form ‘ODE + noise’, which is our main focus here. However, an explicit likelihood is not available for many stochastic process models. Such models are sometimes referred to as ‘simulation-based’, ‘implicit’, or ‘likelihood-free’ (see e.g. [29, 30]). In such cases, we assume an approximate or auxiliary likelihood function [31], also called a synthetic [32, 33] or surrogate [34] likelihood function, is available, which places these simulation-based problems within the proposed framework. In particular, as described by, e.g., [31, 35], also building on the prior work in indirect inference [36, 37], the critical component of approximate likelihood approaches is a function that links the parameters of the model of interest, *θ*, to the parameters *ϕ* that define a tractable probability distribution *p*(*y*; *ϕ*) for the observable data *y*:

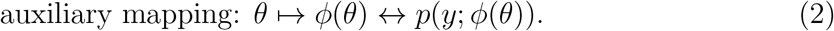

Note that the *↔* notation indicates that this is a bijective mapping, i.e. the data distribution parameters uniquely and minimally define the data distribution. Throughout the manuscript, we use a standard abuse of notation by using the same symbol for both the auxiliary mapping, *ϕ*, and its values, *ϕ*(*θ*), the data distribution parameters. Furthermore, throughout this document all quantities may generally be vector-valued, and we do not use bold for vectors to be consistent with the statistics literature. The data *y* may also represent summary statistics of finer-scale data.

In our framework, the auxiliary function linking mechanistic model parameters to data distribution parameters is conceptually important even in the ‘exact’ cases as it provides a more structured link between the mechanistic model parameters and data distribution parameters than the forward mapping or likelihood alone. The upper left of Figure 1 depicts this role of the auxiliary mapping in determining relationship between the mechanistic model and the auxiliary data distribution model. Hence, although for simplicity we focus on elementary ODE-based models in the present work, in which case the auxiliary mapping leads to an ‘exact’ likelihood, we still make use of an explicit auxiliary mapping for completeness. A very common and familiar example of an auxiliary mapping is a function mapping the parameters of an ODE to the associated trajectory, measured at a discrete set of points, then identified with the mean of a (multivariate) normal distribution for observations. All of our examples in the present manuscript use an auxiliary mapping of this form (e.g. mapping the model parameters to the mean of a normal or Poisson observation distribution), though this is not required in general. In contrast to much of the indirect inference literature [36, 37], here we do not assume the auxiliary mapping is one-to-one, which would correspond to the assumption of an identifiability condition [25]. Instead, we assess this by carrying out an identifiability analysis using the profile likelihood. In the case where auxiliary parameters such as the variance of the noise are unknown, we can include the ‘observational’ model parameters in *θ*, representing the ‘observation mechanism’ [38].

The assumption of requiring an auxiliary likelihood is no real limitation in practice as such synthetic/auxiliary methods typically perform at least as well as alternatives with implicit or nonparametric likelihood functions such as approximate Bayesian computation [31, 32, 34, 35, 39, 40]. Still, we recommend checking approximate-likelihood-based inferences with synthetic data to ensure the approximations preserve the relevant inferential properties, as these are not always guaranteed [41]. With the above discussion in mind, we will generally refer to likelihood functions (specified in detail in Results), simply as ‘likelihood functions’, whether ‘exact’ or approximate, and will not distinguish these further in the present work because we explicitly focus on the ‘exact’ case from this point on.

#### Profile likelihood

The essential tool of our PWA framework is *profile likelihood*, which is obtained from a full likelihood by optimizing out (or eliminating) so-called nuisance parameters. We use this tool in our workflow in both traditional and newly developed ways. Initially established in the conventional statistical literature as a tool for eliminating nuisance parameters and constructing approximate confidence sets with asymptotic guarantees and good performance in small samples [9, 27, 28, 42], profile likelihood is also a reliable and efficient tool for practical identifiability analysis of mechanistic models [13, 43–49, 51, 73]. For example, we have shown that profile likelihood analysis provides equivalent verdicts concerning identifiability orders of magnitudes faster than other approaches, such as Markov chain Monte Carlo (MCMC) [13, 52–54]. Others have shown that the profile likelihood can provide reliable insights when other methods, including MCMC and Fisher information-based methods, may fail [48, 55]. Thus we use profile likelihood as a tool for an initial identifiability analysis in mechanistic models, as proposed by Raue et al. (2013) [48] and our previous work [13]. After this initial identifiability analysis, we also use profile likelihood as an estimation and uncertainty analysis tool, using its good frequentist calibration properties [9, 27]. In addition to efficiently constructing confidence sets representing overall uncertainties, our approach allows us to target our inferences towards specific questions and attribute the influence of specific parameters or combinations of parameters on predictions. We do this by taking advantage of the conventional targeting property of profile likelihood [9], allowing us to form approximate likelihood functions and corresponding confidence sets for so-called *interest parameters*. We then use our recently developed method [22–24] for propagating individual profile-likelihood-based parameter uncertainties through to predictions and associated ‘profile-wise’ confidence sets before recombining these into an overall confidence set accounting for the uncertainties in all parameters. This approach is conceptually simple and computationally straightforward compared to existing approaches as we use combinations of lower-dimensional approximations to the full likelihood function and directly propagate parameter confidence sets forward instead of requiring additional constrained optimization as in the optimal control formulation of prediction profile likelihood [18, 19]. We extend our approach to propagate bivariate profiles for higher accuracy and assess this against the results from the full likelihood. As we show, PWA is useful for targeting specific parameters or combinations of parameters to explore how they impact a predictive quantity of interest. These tools together form a systematic workflow incorporating identifiability analysis, parameter estimation, and prediction, each component capable of targeting particular interest parameters or combinations of parameters, as well as recombined to understand overall uncertainties.

#### Prior information

Like other frequentist and likelihood-based methods, our approach can, in principle, incorporate prior information in the form of prior likelihoods based on past observations [27, 56, 57] and constraints in the form of set constraints [58] without the need for prior probability distributions over parameters [22]. In the present work, we only use basic prior information in the form of set constraints, providing lower and upper bounds on parameters. As discussed by Stark [58], this is distinct from (and makes fewer assumptions than) assuming a uniform prior probability over parameter space as might be done in a Bayesian analysis. We mention this to emphasize that in our view, the ability to incorporate past information is not a core distinguishing feature of Bayesian and frequentist analyses (see also [56] and [58]), as opposed to their distinct goals and interpretations [8, 9].

#### Identifiability analysis and uncertainty analysis: full likelihood methods

Given a likelihood function linking our mechanistic model parameters to data via the auxiliary mapping, we can carry out practical identifiability (or estimability [25]) analysis and statistical inference. As indicated above, we take a likelihood-based frequentist approach to statistical inference [9, 27, 28, 42]. We consider the likelihood function as an intuitive measure of the information in the data about the parameters and construct approximately calibrated confidence sets for parameters, *C*_*θ*_, by thresholding the likelihood function according to standard sampling theory [27]. In addition to providing confidence sets for the whole parameter vector with good frequentist (i.e., repeated sampling) properties for sufficiently regular problems, we can also use the likelihood to diagnose and analyse structurally or practically non-identifiable models using profile likelihood [13, 18, 19, 47], as well as to construct approximate, *targeted* confidence intervals for interest parameters [9, 27, 42].

While likelihood and profile likelihood are commonly used to estimate and analyse information in the data about mechanistic model *parameters* [13, 47], *they have been less widely used in this context for predictions* (though see [18, 19, 22–24]). However, likelihood-based confidence set tools are well-suited for this purpose. Firstly, one can, in principle, directly propagate a likelihood-based confidence set for parameters, *C*_*θ*_, forward under the auxiliary mapping *ϕ* from mechanistic model parameters to data distribution parameters, to construct a confidence set, *C*_*ϕ*_, for the data distribution parameters. For example, a confidence set for the parameters *θ* of an ODE, d*z/*d*t* = *f* (*z*; *θ*), can be propagated under the model to its solution *z*(*θ*). We might then define the auxiliary mapping such that the ODE solution represents the mean of the data distribution, *p*(*y*; *ϕ*(*θ*)) = *p*(*y*; *z*(*θ*)) for density *p*(*y*; *ϕ*) with mean parameter *ϕ*. This process, illustrated in the centre panel in Figure 1, allows us to form a confidence set for *ϕ* and is described in detail in the Results section. As also shown in the figure, we can further construct confidence sets *C*_*y*_ for (noisy) *data realisations*, which leads to additional uncertainty. The term prediction interval is often reserved for such predictions of realisations in statistical applications [9, 59, 60] while here, because of the mechanistic interpretation of the data distribution parameters (including the ability to project out of sample), we refer to both the data distribution parameters *ϕ*(*θ*) and realisations *y*, drawn from the associated distribution *p*(*y*; *ϕ*(*θ*)), as ‘predictive’ quantities.

#### Identifiability analysis and uncertainty analysis: profile-wise profile likelihood methods

The above full likelihood approach is conceptually simple, but can be computationally expensive or infeasible when the parameter dimension is large. Furthermore, while this approach simultaneously accounts for uncertainties in all parameters, it makes it difficult to determine how uncertain our estimates of particular parameters, or combinations of parameters, are and to attribute different contributions to the total uncertainty in predictions from specific parameters or combinations of parameters. These factors make profile likelihood an attractive approach to approximately target parameters and investigate their influence on predictive quantities. The latter usage – a form of approximate statistical sensitivity study linking mechanistic and data distribution parameters and providing associated confidence sets – is something we developed recently [22–24]. A similar idea was independently introduced in the context of local derivative-based sensitivity analysis [61, 62], where the authors consider the total derivative of outputs with respect to an interest parameter along the associated profile curve. While similar in spirit, our approach is fully nonlinear, global over the profiling range, can generalise to vector interest parameters, and provides approximate confidence sets for outputs rather than sensitivities expressed as derivatives.

The left of the bottom row of Figure 1 shows how we can approximately decompose the overall parameter uncertainty into components based on profile likelihoods. Importantly, these components are efficiently computable separately and can be propagated separately before being re-combined to approximate the total uncertainty in predictive quantities. This decomposed approach forms the basis of the PWA workflow for estimation and prediction with mechanistic mathematical models presented in this work. In addition to approximating the more computationally-expensive, complete likelihood methods, our approach allows for additional insight into the *attribution* of uncertainty by *targeting* our inferences and propagation of predictions.

### Manuscript structure

This manuscript is structured as follows. As our main contribution is a new methodology, we present the details of our approach in the Results section. In Results, we first provide the details of workflow, methodology and notation alongside a gold-standard full likelihood approach. We then present three case studies focusing on canonical ODE-based models used in population biology and ecology applications, together with Gaussian and non-Gaussian error models. Our case studies emphasise how to use parameter confidence sets to construct approximate profile-wise and combined confidence sets for predictive quantities, as this predictive component of likelihood-based methodologies is the least explored in the literature. This component also completes a natural analysis cycle, from data to model parameters and back to predictions, enabling an iterative approach to model building, inference, and checking. Finally, in the ‘Conclusions and future directions’ section, we summarise our contribution and mention areas for future consideration. We note that all computational examples can be replicated using software written in the open-source Julia language made available on GitHub.

## Materials and methods

Since this study describes a new computational methodology, we present the mathematical details for the components of the methods in the Results section.

## Results

Here we provide full details of the core components of the workflow illustrated in Figure 1, before considering a series of case studies. We describe our workflow for dynamic, differential equation-based models for simplicity. However, the same ideas carry over to other models, e.g., stochastic differential equations [63], spatiotemporal models such as partial differential equations (PDEs) [13] or simulation-based stochastic models [35].

### Full likelihood methods

We first define the probability model, data, and full likelihood function. As mentioned, our focus here is on dynamic mechanistic models – ODE models in particular – so we specify our workflow components in these terms. Again as mentioned, the generalisation to other cases is straightforward conceptually. We also explicitly layout the logic of frequentist uncertainty propagation in the subsection ‘Likelihood-based confidence sets for data distribution parameters’, and hence justify the classification of this ‘full likelihood’ approach to prediction as a gold standard method.

### Observed data

We assume that observed data *y*° are measured at *I* discrete times, *t*_*i*_, for *i* = 1, 2, 3, …, *I*. When collected into a vector, we denote the noisy data by 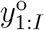, where the subscript makes it clear that the data consists of a time series of *I* observations. Multiple measurements at a single time can be represented by repeated *t*_*i*_ values for distinct *i* indices.

### Mechanistic model

The mechanistic models of interest here are assumed to satisfy differential equations, i.e., of the general form

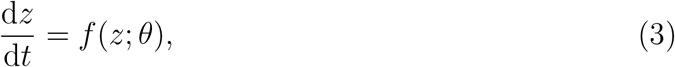

where *θ* is the vector of mechanistic model parameters and *z* is also vector-valued in general. We denote noise-free model solutions evaluated at observation locations *t*_*i*_ by *z*_*i*_(*θ*) = *z*(*t*_*i*_; *θ*). As with the noisy observations, we denote the process model solution by *z*_1:*I*_(*θ*) for the model solution vector at the discrete time points and by *z*(*θ*) for the full (continuous) model trajectory over the time interval of interest.

### Auxiliary mapping

The mapping from model parameters to data distribution parameters takes the general form:

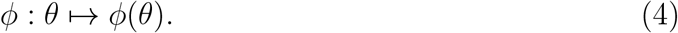

Here we will assume the model solution (trajectory) is the key data distribution parameter. In particular, we take the model solution to correspond to the mean of the data distribution. Hence we have

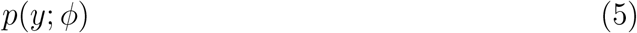

for density *p* with mean parameter *ϕ* and the mapping

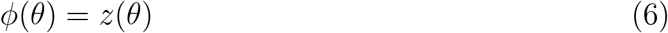

defining the data mean as equal to the model solution. At observation locations *t*_*i*_ we will define

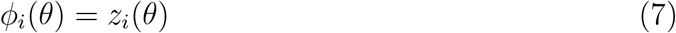

where *z*_*i*_(*θ*) is defined above.

As discussed below, we will assume the variance in normal distribution data models is known for simplicity, though we can estimate this simultaneously [38]. We also consider the case of Poisson noise, for which the mean defines the variance.

### Likelihood function

Given the above elements, we have a density function for the observable random variable *y* depending on the model parameter *θ* of the form

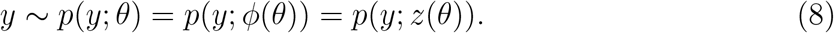

At any given observation time, *t*_*i*_, we have

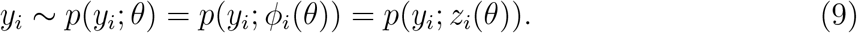

For a collection of observations 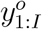, treating the distribution of the observations as a function of the parameter *θ* with observations fixed, we can define the normalised likelihood function as [28]

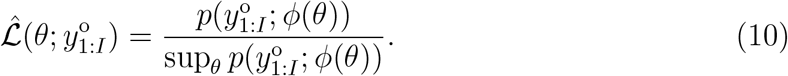

Often the likelihood is only defined up to a constant, but here we fix the proportionality constant by working with the normalised likelihood function. The normalised likelihood function is also appropriate for forming asymptotic confidence intervals [27]. The hat notation indicates the normalisation – when required, the unnormalised likelihood function (and unnormalised log-likelihood) will be denoted without the hat. We will occasionally refer to the likelihood function as the ‘full’ likelihood function to distinguish it from the profile likelihood functions considered below.

We will assume that the observations at each time are independent of the others given the model trajectory at that time, i.e.

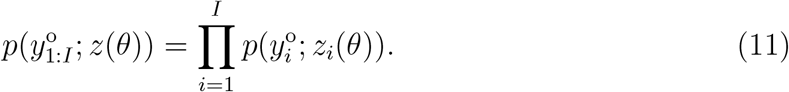

In practice, we work with the log-likelihood function, which, given a set of *I* independent (given the model trajectory) observations, takes the form

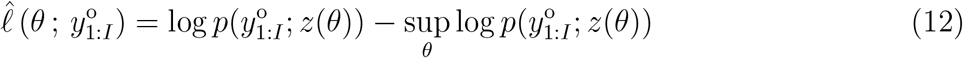

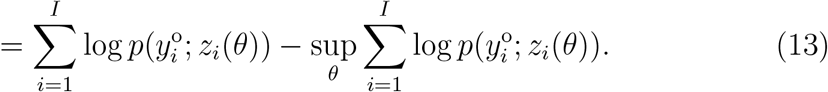

The normalising factor, i.e. the value associated with the maximum likelihood estimate (MLE) 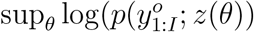, can be computed by numerical optimization of the unnormalised log-likelihood function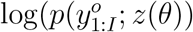. In the case of normally-distributed noise about the model trajectory *z*(*θ*), with known variance *σ*^2^, we have the unnormalised log-likelihood function

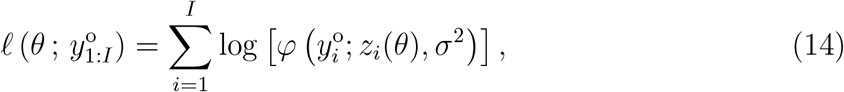

where *φ*(*x*; *µ, σ*^2^) denotes a Gaussian probability density function with mean *µ* and variance *σ*^2^, and *ϕ*(*θ*) = *z*(*θ*) defines the auxiliary mapping from model parameters to the mean of the normal distribution. Similar expressions hold for non-normal models; we discuss the case of a Poisson noise model in one of our case studies below.

### Likelihood-based confidence sets for parameters

Given the log-likelihood function, we can form approximate likelihood-based confidence sets for *θ* using [27, 28]

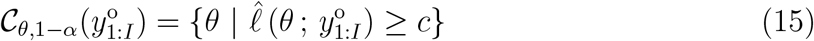

for a cutoff parameter, *c*, chosen such that the confidence interval has approximate coverage of 1 *− α* (the usual choice being 95% i.e. *α* = 0.05). This threshold is, conceptually, based on inverting a likelihood ratio test for each candidate parameter value and can be calibrated for sufficiently regular problems using the chi-square distribution, *ℓ*^*∗*^ = *−*∆_*ν,q*_*/*2, where ∆_*ν,q*_ refers to the *q*th quantile of a *χ*^2^ distribution with *ν* degrees of freedom (taken to be equal to the dimension of the parameter) [27]. This calibration is with respect to the (hypothetical) ‘repeated sampling’ of *y*_1:*I*_ under the fixed, true, but unknown distribution [9]. Above, 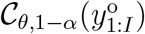 is the realised confidence set from the random variable *C*_*θ*,1*−α*_(*y*_1:*I*_) where *y*_1:*I*_ *∼ p*(*y*_1:*I*_; *θ*). We will drop the explicit dependence on the random variable *y*_1:*I*_ in general, but we emphasise that all probability statements in the frequentist approach are made under the distribution of *y*_1:*I*_.

### Likelihood-based confidence sets for data distribution parameters

Given a confidence set for the model parameters and the auxiliary mapping *ϕ*(*θ*), we can define a corresponding confidence set *𝒞*_*ϕ*_ for the data distribution parameters *ϕ*. In particular, we define this confidence set as the image of the parameter confidence set under the auxiliary mapping *ϕ*:

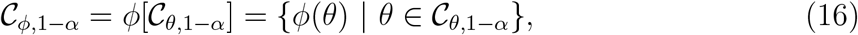

where the square brackets indicate that we are taking the set image under the mapping. Since the definition above implies that we have the following relationship between statements:

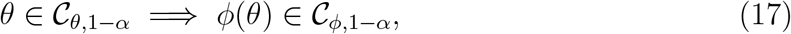

it follows that the confidence set *𝒞*_*ϕ*,1*−α*_ has *at least* 95% coverage (i.e. is conservative). The intuitive reason for this is that when the above implication statement holds, and under repeated sampling, the statements of the form *ϕ*(*θ*) *∈ 𝒞*_*ϕ*,1*−α*_ are true in all cases that the statements *θ ∈ 𝒞*_*θ*,1*−α*_ are true. So the proportion of correct statements about *ϕ* under repeated sampling must be at least as high as those made for *θ*. There may, however, be cases in which *θ ?∈ 𝒞*_*θ*,1*−α*_ yet *ϕ*(*θ*) *∈ 𝒞*_*ϕ*,1*−α*_, and hence the coverage for our data distribution parameters is at least as good as for our parameters – we may occasionally correctly predict *ϕ* using the wrong model parameter. If the mapping *ϕ* is 1-1, the statements are equivalent, and the two confidence sets will have identical coverage.

We classify this full likelihood approach as gold-standard for likelihood-based methods as, from the above arguments, the usual guarantees for likelihood-based model parameter confidence sets directly translate to guarantees for data-distribution (predictive) parameters. We also note that, in principle, this approach provides an alternative way to compute the usual prediction profile likelihood, as given in [19], since a complete grid evaluation of the likelihood will, by definition, determine the highest likelihood value associated with the prediction of interest. In practice, when dealing with the full likelihood, we must resort to Monte Carlo or other related approximate/ensemble methods [20], e.g. evaluating and propagating a finite set of parameter vectors meeting the full likelihood cutoff. Nevertheless, even with computational approximations, we view this approach as the natural standard to assess our profile-wise approximation against.

In our approach we generate confidence sets for the predictive quantity *ϕ*(*θ*) by first forming confidence sets for the parameter *θ*. Here, the mechanistic model provides a natural constraint indexed by a finite-dimensional parameter. Usefully, this means we can form confidence sets for ‘infinite-dimensional’ quantities, such as full model trajectories, based on finite-dimensional confidence sets for the parameters. This ability is another advantage of mechanistic-model-based inference – forming confidence sets for functions/curves is a difficult problem without additional constraints, and fixed-time (i.e., pointwise) intervals tend to underestimate extremes in dynamic models [64]. Furthermore, our (full likelihood) confidence sets have coverage guarantees for data distribution parameters that are at least as good as those for the parameters, in contrast to confidence sets based on linearisations, and have non-constant widths in general.

### Likelihood-based confidence sets for data realisations

Our primary focus in the present work is on a workflow involving confidence sets for mechanistic model parameters and data distribution parameters. Under ‘data distribution parameters’, we include quantities such as the model trajectory considered as the data mean. We call the latter a ‘predictive’ quantity in the context of mechanistic models as it is the natural output of simulating or solving the model, and it can be projected forward beyond the data, providing a prediction of a future data average. However, the term ‘prediction’ is often used when discussing predictions of data realisations (including e.g. measurement noise), i.e., for *y ∼ p*(*y*; *ϕ*) rather than for distribution parameters *ϕ* [9, 59, 60]. These noisy realisations are another type of ‘predictive’ quantity of interest for which we can form confidence sets, though the process is, in general, more complicated. This is because the error rate when predicting a noisy realisation from a data distribution with estimated parameter depends on both the error rate in estimating the parameter and the inherent error involved in predicting a stochastic quantity. For completeness, we sketch the process of forming such sets, and how to obtain bounds on the associated combined error rates, but do not go into detail. Figure 1 also illustrates this. Readers who are interested in estimating the model mean trajectory only can skip this section.

To see how to form a prediction set for noisy realisations, first, consider predicting a single unseen *y*_*j*_ given full knowledge of *p*(*y*; *ϕ*) and assuming *y*_*j*_ *∼ p*(*y*; *ϕ*). Given exact knowledge of *ϕ* this task is simple. By definition, we need to identify, for the given *ϕ*, a subset of the sample space *𝒜*_1*−α*_(*ϕ*) having probability 1 *− α*. This is trivial for simple parametric models such as the normal distribution where one can take a region of plus or minus approximately two standard deviations. The main difficulty is that, for our application of interest, we only have an uncertain estimate of (i.e., confidence set for) *ϕ*. In addition, this set is not of a known form in general, as we do not use linearisation to propagate our parameter confidence sets. Under complex conditions such as these, a simple, general, conservative approach is to use a Bonferroni correction method (such methods are common in the simultaneous inference literature [65]): we form a confidence set *𝒞*_*ϕ*,1*−α/*2_ for *ϕ* at the level 1 *− α/*2, and, for each *ϕ*, construct an appropriate subset *𝒜*_1*−α/*2_(*ϕ*) such that it has probability 1 *− α/*2 of correct prediction for the distribution indexed by that *ϕ*, where ‘correct prediction’ means *y*_*j*_ *∈ 𝒜*_1*−α/*2_(*ϕ*). Taking the union of the resulting sets, i.e.

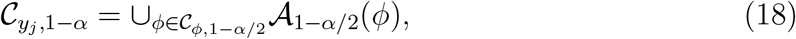

gives a conservative confidence set for *y*_*j*_ at the level 1 *− α* for realisations *y*_*j*_. Intuitively, the error rate *α* is (bounded by) the sum of the two sources of error: the data distribution parameter and the data realisation from the associated distribution. Thus we need to form confidence sets at a higher confidence level (e.g., 97.5%) for each component quantity to ensure the resulting coverage for our goal (e.g., 95%).

Though the proof that this is a valid (though conservative) prediction interval is trivial, and the same basic idea appears in the literature on constructing ‘tolerance intervals’ [65, 66], we are unaware of a standard reference in the literature for this type of usage. While other approaches tend to target the predictions directly [59, 60], our approach utilises the availability of confidence sets for parameters of data distributions. The same procedure can also be extended to the simultaneous prediction of multiple future observations, though, as with other applications of Bonferroni corrections [65], these may become even more conservative.

### Profile-wise methods

The section ‘Full likelihood methods’ (above) outlines the basic approach to forming and propagating confidence sets based on the full likelihood function associated with a mechanistic model and an auxiliary mapping connecting the model parameters to the parameters of a data distribution. We now consider using profile likelihood methods for both targeting and improving the computational efficiency of our inferences. In this section, we work with log-likelihood functions exclusively, as these are how we implement our methods in practice.

#### Profile likelihood function

To assess the practical identifiability of our problem and to target our inferences, we take a profile likelihood-based approach to inspect 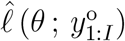 in terms of a series of simpler functions of lower dimensional interest parameters. For simplicity, we assume the full parameter *θ* is partitioned into an interest parameter *ψ* and a nuisance parameter *ω*, such that *θ* = (*ψ, ω*). In general, an interest parameter can be defined as any function of the full parameter vector *ψ* = *ψ*(*θ*), and the nuisance parameter is the complement, allowing the full parameter vector to be reconstructed [42, 67]. For a set of data *y*° the profile log-likelihood for the interest parameter *ψ* given the partition (*ψ, ω*) is defined as [9, 27]

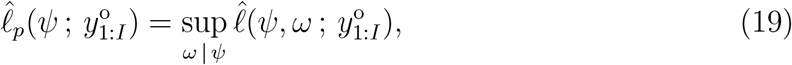

which implicitly defines a function *ω*^*∗*^ (*ψ*) of optimal values of *ω* for each value of *ψ*, and defines a surface with points (*ψ, ω*^*∗*^(*ψ*)) in parameter space. The bottom left of Figure 1 shows profile likelihood functions for parameters. For convenience, Figure 1 shows profiles for both interest and nuisance parameters, *ψ* and *ω*, respectively, but more precisely, we take each parameter as the interest parameter in turn. This series of optimisation problems can be solved using the same algorithm as used for computing the MLE (though typically with a better starting guess, e.g. the MLE or previous point on the profile curve).

#### Profile likelihood-based confidence sets for parameters

Approximate profile likelihood-based confidence sets are formed by applying the same dimension-dependent chi-square calibration, taking the dimension as the interest parameter’s dimension [27]. As for the full likelihood, this again leads to approximate confidence sets, but now for interest parameters, i.e., our inferences are targeted.

#### Profile likelihood-based confidence sets for data distribution parameters

There are at least two obvious ways to construct confidence sets for predictive quantities, such as the model (mean) trajectory. Firstly, we can follow the approach outlined in the section on full likelihood methods above, propagating parameter confidence sets forward. Secondly, we can directly consider profile likelihood for predictive quantities defined as a parameter function. However, both of these approaches present computational difficulties and interpretational limitations. The first requires determining a potentially high-dimensional confidence set from the full likelihood function; the second requires solving optimisation problems subject to constraints on predictions; for example, the profile likelihood for the model trajectory *z*(*θ*) requires solving an optimisation problem subject to a series of trajectory constraints. In the latter case, the literature typically focuses on predictions at a small number of time points [18, 19, 22], which is easier to implement computationally.

We recently introduced an alternative approach to producing approximate profile predictions [22, 23, 38]. We called these parameter-wise profile predictions previously; to emphasise that these can also be based on bivariate or higher-order profiles, here we call these *parameter-based, profile-wise predictions* or *profile-wise predictions* for short. This approach is easier to compute with but also provides a way of approximately *attributing* the uncertainty in predictions to particular parameters or combinations of parameters.

Our approach works by using a series of profile likelihoods and associated confidence sets for *parameters*, then propagating these forward to *predictions*. These individual confidence sets for predictions provide an indication of the influence of particular parameters or combinations of parameters. These individual confidence sets can then be combined to form an approximate confidence set representing the combined contributions to the uncertainty in the predictive quantity of interest. Our approach to confidence sets for predictive quantities is shown in the bottom row of Figure 1, the second two columns.

We can compute this profile likelihood for *ϕ* as follows. We assume a partition of the full parameter into an interest parameter *ψ* and nuisance parameter *ω* (both of which may be vector-valued) for simplicity. This computation produces a curve or hyper-surface of values (*ψ, ω*^*∗*^(*ψ*)). The idea of our approach is then straightforward: we substitute this set of parameters (*ψ, ω*^*∗*^(*ψ*)) into our predictive quantity of interest, here data distribution parameters *ϕ*, i.e., we compute *ϕ*(*ψ, ω*^*∗*^(*ψ*)). The approximate ‘profile-wise likelihood’ for any given *ϕ*, on the basis of *ϕ*(*ψ, ω*^*∗*^(*ψ*)), is defined by taking the maximum (sup) profile likelihood over all *ψ* values consistent with *ϕ*, i.e.

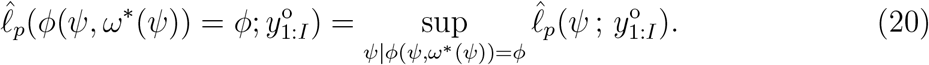

This definition follows the process of constructing a profile for any well-defined function *ϕ* of *θ* using a standard likelihood in *θ*, but now starting from a profile likelihood based on *ψ*. If *ϕ*(*ψ, ω*^*∗*^(*ψ*)) is 1-1 in *ψ* then this is simply the profile likelihood value of *ψ*, but this is not a necessary condition. Computation is carried out by first evaluating *ϕ* along the profile parameter curve/set (*ψ, ω*^*∗*^(*ψ*)) and assigning it tentative profile likelihood values equal to the likelihood value of (*ψ, ω*^*∗*^(*ψ*)). The final profile likelihood value is then taken as the maximum value over compatible *ψ* values. The approach is simple and computationally cheap – for a one-dimensional interest parameter we only need to evaluate the predictive quantity along a one-dimensional curve embedded in parameter space. Here we also consider two-dimensional profile-wise predictions.

Our profile-wise prediction tool enables the construction of approximate intervals for predictive quantities when the parameter of interest (scalar or vector-valued) is considered the primary influence. Consequently, this can illustrate which aspects of a prediction depend most strongly on the interest parameter (e.g., the early or later parts of the mean trajectory). While desirable for understanding the impacts of individual parameters on predictions, this feature means the particular intervals will typically have lower coverage for the prediction than other approaches taking into account all uncertainties simultaneously (due to neglected dependencies on the nuisance parameters). However, we can reconstruct an approximate overall predictive confidence set accounting (to some extent) for all parameters by combining the individual profile-wise prediction sets to give a combined confidence set. Our approach aims to approximate what would be obtained by propagating the parameter confidence sets formed from the full likelihood function. In particular, the definition of coverage for a confidence set means that, given any collection of confidence sets for the same quantity, more conservative confidence sets for that quantity can always be constructed by taking the union over all sets. Thus we take the set *union* of our individual prediction sets:

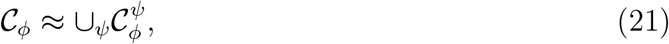

for a series of interest parameters *ψ* (typically chosen to be each parameter component, or all pairs of parameter components, of the full parameter in turn). Furthermore, by increasing the dimension of the interest parameter, we obtain a better approximation to the prediction set based on the full likelihood. An open question is to what extent lower-dimensional approximations can be trusted, and we continue our investigations of this here. We have previously applied unions of one-dimensional profiles [23, 24]; here, we extend this to unions of two-dimensional profiles in addition to unions of one-dimensional profiles.

#### Profile likelihood-based confidence sets for realisations

Given an approximate confidence set for *ϕ*, formed according to our approximate profile likelihood approach, and the goal of producing a confidence set for an unseen *y*_*j*_, we can proceed exactly as in the full likelihood case (see Figure 1 far right of bottom row).

We now consider three case studies to showcase the workflow. In all cases, we work with ODE-based mechanistic models since these types of models exemplify the kinds of commonly deployed models for a range of important applications, for example, biological population dynamics [68], disease transmission [69] and systems biology [70], among many others. The first case study is a simple single nonlinear ODE model with Gaussian noise, and the two remaining case studies seek to introduce important complexities. The second case study works with a system of coupled nonlinear ODEs with Gaussian noise, while the third and final case study focuses on a single nonlinear ODE model with a non-Gaussian noise model. We provide software, written in Julia, on GitHub to replicate these results and to facilitate applying these ideas to other mathematical and noise models. Note that we deliberately focus the presentation of the workflow on identifiable and reasonably well-behaved problems which means that we are able to use standard, local search-based numerical optimization tools with default stopping criteria. In more complex problems, more sophisticated optimization tools may be required to avoid local optima.

### Case Study 1: Logistic growth with Gaussian noise

We now demonstrate our workflow starting with the canonical logistic growth model that is used throughout biology and ecology to describe sigmoid growth of a population with near exponential growth at small population densities and zero net growth as the population density approaches a carrying capacity density [71]. The logistic growth model has applications covering a phenomenal range of length and time scales ranging from microscopic growth of cancer cell populations [23] to reef-scale growth of corals on coral reefs [38] to continental-scale growth of human populations [72]. The dimensional logistic model assumes that the population density *C*(*t*) *≥* 0 evolves according to

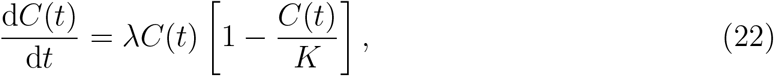

where *λ >* 0 is the intrinsic growth rate and *K* is the carrying capacity density. To solve Eq (22) we need to specify the initial density, *C*(0). Therefore, the parameter vector for this model is *θ* = (*λ, K, C*(0)). While the logistic model has an exact solution, we aim to keep our workflow as general as possible so we solve all models in this work numerically [74]. For a Gaussian noise model we have

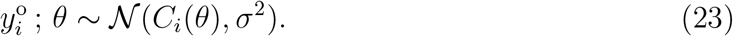

Here we take *σ* as a pre-specified constant, which is often used in practice [13, 70], but *σ* can also be estimated with the other parameters, if required [38]. Data in Figure 2(a) shows 11 equally-spaced noisy observations on the interval 0 *≤ t ≤* 1000 superimposed with the MLE solution of Eq (22). All calculations in this work that involve numerical optimisation are obtained using the Nelder-Mead algorithm with simple bound constraints [75].

**Figure 2:**
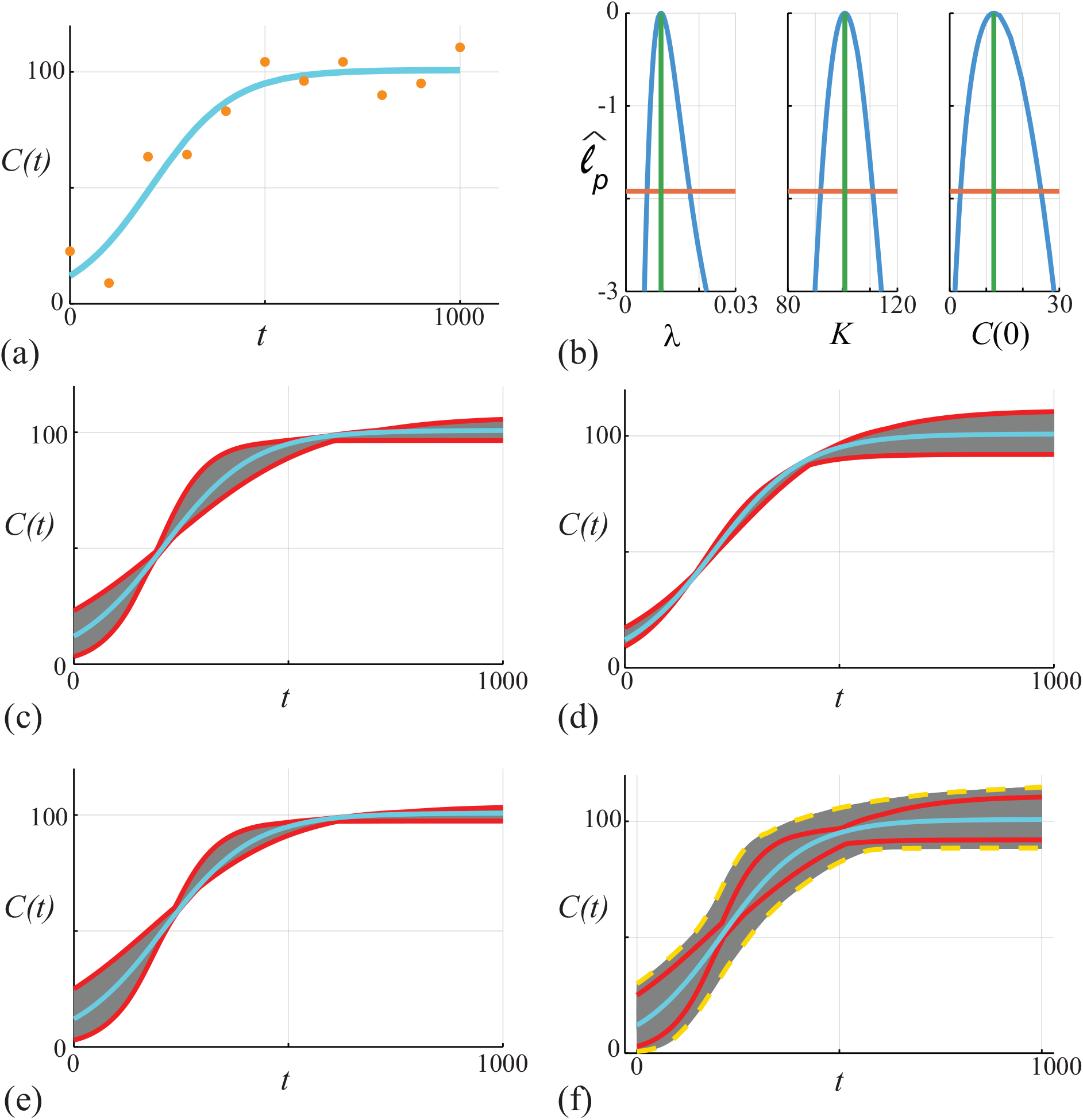
(a) Data obtained by solving Eq (22), with *θ* = (*λ, K, C*(0)) = (0.01, 100, 10), at *t* = 0, 100, 200, …, 1000 is corrupted with Gaussian noise with *σ* = 10. The MLE (cyan) solution is superimposed, 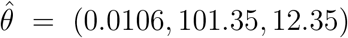. (b) Univariate profiles for *λ, K* and *C*(0), respectively. Each profile is superimposed with the MLE (vertical green) and the 95% threshold 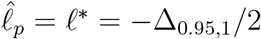 is shown (horizontal orange). Points of intersection of each profile and the threshold define asymptotic confidence intervals: 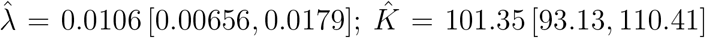 and, 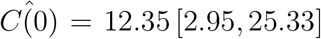. (c)-(e) *C*(*t*) trajectories associated with the *λ, K* and *C*(0) univariate profiles in (b), respectively. In each case we uniformly discretise the interest parameter using *N* = 100 points along the 95% confidence interval identified in (b), and solve the model forward with *θ* = (*ψ, ω*) and plot the family of solutions (grey), superimposing the MLE (cyan), and we identify the prediction interval defined by these solutions (solid red). (f) Compares approximate prediction intervals obtained by computing the union of the univariate profiles with the ‘exact’ prediction intervals computed from the full likelihood. Trajectories (grey) are constructed by considering *N* = 10^4^ choices of *θ* and we plot solutions with 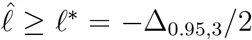 only. These solutions define an ‘exact’ (or gold-standard) prediction interval (dashed gold) that we compare with the MLE solution (cyan) and with the approximate prediction interval formed by taking the union of the three univariate intervals (red).

To assess the practical identifiability of this problem we calculate univariate profile likelihood functions shown in Figure 2(b). As for calculating the MLE, univariate profile likelihoods are obtained using the Nelder-Mead algorithm [75]. For this model we have three univariate profile likelihoods obtained by considering: (i) *ψ* = *λ* and *ω* = (*K, C*(0)); (ii) *ψ* = *K* and *ω* = (*λ, C*(0)); and, (iii) *ψ* = *C*(0) and *ω* = (*λ, K*), respectively. In each case the univariate profiles are well formed about a single distinct peak, and we form a confidence interval by estimating the interval where 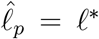. We use the bisection algorithm to determine the point(s) of intersection. In the case of *ψ* = *λ* we determine the two points of intersection, *λ*_min_ and *λ*_max_.

Before considering prediction we first explore the empirical, finite sample coverage properties for our parameter confidence intervals as they are only expected to hold asymptotically [27]. We generate 5000 data realisations in the same way that we generated the single data realisation in Figure 2(a), for the same fixed true parameters. For each realisation we compute the MLE and count the proportion of realisations with 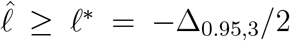, where 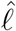 is evaluated at the true parameter values (this condition means the true parameter vector is within the sample-dependent, likelihood-based confidence set). This gives 4753*/*5000 = 95.06%, which is very close to the expected asymptotic result. In addition to checking coverage of the confidence sets based on the full likelihood function, for each of the 5000 data realisations we separately compute the univariate profiles for *λ, K* and *C*(0). For each univariate profile we check whether the 95% confidence interval of that profile contains the true value, which we find holds for 94.74%, 94.95% and 94.70% of realisations for *λ, K* and *C*(0), respectively [73].

For each univariate profile in Figure 2(b) we construct a profile-wise prediction interval for the density, *C*(*t*) [24]. For example, to interpret how uncertainty in *λ* translates into uncertainty in *C*(*t*) we uniformly discretise the interval (*λ*_min_, *λ*_max_) into *N* points and use numerical optimisation to compute the nuisance parameters at each point. For each of the *N* parameter combinations, we solve Eq (22) and plot the family of solutions in Figure 2(c) (grey solid curves). This family of solutions allows us to construct a prediction interval for the model trajectory (red solid curves). The profile-wise prediction interval for *λ* leads to a relatively wide prediction interval during early time as *C*(*t*) increases, whereas the prediction interval narrows at later time. This is intuitively reasonable since *λ* does not impact the late time solution of Eq (22). Profile-wise prediction intervals for *K* and *C*(0) are also constructed, as shown in Figure 2(d) and (e), respectively. Again, these prediction intervals provide clear and intuitive insight into how uncertainty in each parameter affects the predictions in different ways. For example, Figure 2(d) shows a relatively wide prediction interval at late time, which is again intuitively obvious since the late time solution of Eq (22) depends solely on the value of *K*. Similarly, Figure 2(e) shows a relatively wide prediction interval at early time, which also intuitively reasonable since the early time solution depends on the estimate of *C*(0). Based on these three profile-wise prediction intervals we then compute the union of the intervals, shown in Figure 2(f), where it is clear that the union encloses the MLE solution.

To explore the accuracy of the union of profile-wise prediction intervals we compare these results with prediction intervals obtained by working with the full likelihood function. We work with the full likelihood function by randomly sampling *N* = 10^5^ locations in the (*λ, K, C*(0)) parameter space and computing 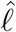 for each choice. For locations where 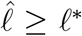 we solve Eq (22) and plot these solutions in Figure 2(f) (grey solid curves). Using these solutions we define a prediction interval (yellow solid curves), and numerical experimentation shows that when *N* is sufficiently large this prediction interval is relatively insensitive to *N*. Comparing the two sets of prediction intervals in Figure 2(f) indicates that both are approximately symmetric about the MLE, however the prediction interval based on the full likelihood (yellow solid curves) is noticeably wider than the approximate prediction interval obtained by taking the union of univariate profiles (red dashed curves) [24]. Despite the difference in prediction intervals it is relevant to note the difference in computational effort required to compute the prediction intervals. Predictions from each univariate profile involves solving the mathematical model forward 10^2^ times, which means that the union of three univariate profiles involves solving the mathematical model forward 3 *×* 10^2^ times since we have three parameters. In contrast, working with the full likelihood model involves solving the mathematical model forward 10^5^ times, which translates into an order of magnitude difference in the computational time to form these two prediction intervals. The difference in prediction intervals between the union of univariate prediction intervals and the prediction intervals obtained with the full likelihood motivates us to consider an improved approximation based on constricting pairwise parameter predictions.

Results in Figure 3(a) show the same data superimposed on the same MLE solution of Eq (22) as reported previously in Figure 2(a). We proceed by computing the three bivariate profiles by setting: (i) *ψ* = (*λ, K*) and *ω* = *C*(0); (ii) *ψ* = (*λ, C*(0)) and *ω* = *K*; and, (iii) *ψ* = (*K, C*(0)) and *ω* = *λ*, respectively. As a concrete example, consider the bivariate profile likelihood for (*λ, K*), given by

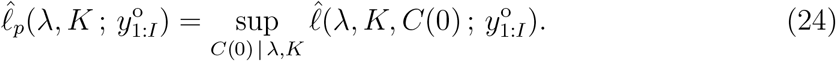

Again, as for calculating the MLE and the univariate profile likelihoods, we calculate all bivariate profile likelihood functions using the Nelder-Mead algorithm [75]. There are many options for how we calculate the bivariate profile and here we simply estimate each bivariate profile on a uniform 20 *×* 20 mesh and superimpose the 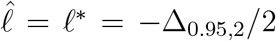 contour (solid red curve). Our initial estimate for the iterative solver at each mesh point is taken to be the MLE for simplicity. The left-most panels in Figure 3(b)-(d) shows the uniform meshes on which we compute the (*λ, K*), (*λ, C*(0) and (*K, C*(0)) bivariate profiles, the associated 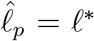 contour, and the MLE (pink disc). For each mesh point within the contour we solve Eq (22) and plot the resulting family of solutions in the right-most panels of Figures3(b)-(d) to form a pairwise family of prediction intervals (solid grey curves). The shape of the 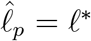 contours provides insight into the pairwise roles of the parameters. The banana-shaped contour in Figure 3(c) suggests that *λ* and *C*(0) are related in terms of their effect on the model solution, which is intuitively reasonable since smaller values of *C*(0) require larger values of *λ* for the solution of Eq to match the data. Similarly, the relatively elliptical shape of the contour in Figure 3(d) indicates little relation between *C*(0) and *K* in terms of their effect on the model solution, which is also reasonable since these quantities are not mechanistically linked in the same way as *C*(0) and *λ* are. Taking the union of the pairwise prediction intervals in Figure 3(e) leads to a prediction interval (dashed red curves) that provides a closer match to the prediction interval formed by working with the full likelihood function (yellow solid curves). Despite this improvement over the union of univariate profiles, the banana shaped contour in Figure 3(c) clearly requires a fine discretisation of the bivariate profile to accurately resolve the shape of the boundary which motivates a second method of dealing with predictions from bivariate profiles.

**Figure 3:**
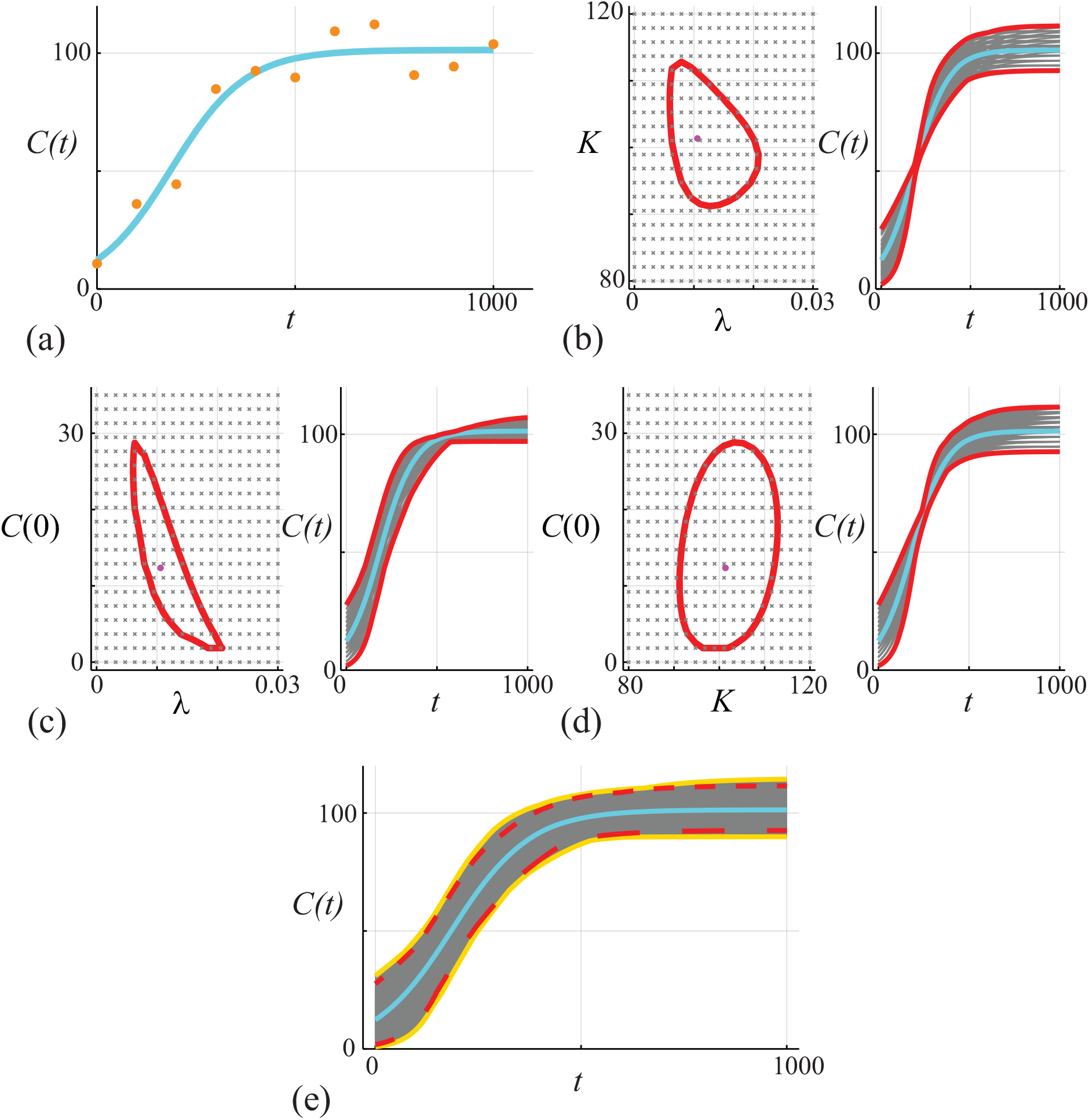
(a) Data obtained by solving Eq (22), with *θ* = (*λ, K, C*(0)) = (0.01, 100, 10), at *t* = 0, 100, 200, …, 1000 is corrupted with Gaussian noise with *σ* = 10. The MLE solution (cyan) is superimposed, 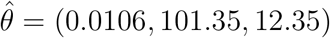. (b)-(d) Bivariate profiles for (*λ, K*), (*λ, C*(0)) and (*C*(0), *K*), respectively. In (b)-(d) the left-most panel shows the bivariate profile uniformly discretised on a 20 × 20 grid with the 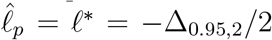 contour and the MLE (pink disc) superimposed. The right-most panels in (b)-(d) show *C*(*t*) predictions (grey) for each grid point contained within the 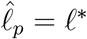 contour, together with the profile-wise prediction interval (solid red). (e) Compares approximate prediction intervals obtained by computing the union of the profile-wise prediction intervals with the ‘exact’ (more precisely, conservative, assuming the parameter confidence set has proper coverage) prediction intervals from the full likelihood function. Predictions (grey) are constructed by considering *N* = 10^4^ choices of *θ* and we plot solutions with 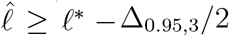 only. These solutions define a gold-standard prediction interval (dashed gold) that we compare with the MLE solution (cyan) and with the approximate prediction interval formed by the union of the three bivariate trajectories (red).

We now introduce a more targeted means of using the bivariate profiles to construct approximate prediction intervals using data in Figure 4(a), which are the same observations and MLE solution of Eq (22) as in Figures 2(a)–3(a). We consider the same three bivariate profiles as in Figure 3 except that for each bivariate profile we target points on the 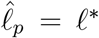 boundary rather than naively meshing the bivariate parameter space. Consider the (*λ, K*) bivariate profile in Figure 4(b), we identify points on the boundary using the following simple algorithm:

1. Randomly choose a value of the abscissa by sampling *λ ∼* U(*λ*_min_, *λ*_max_), where *λ*_min_ and *λ*_max_ are suitable lower and upper bounds (e.g. these bounds could be the same bounds used to compute the MLE).
2. Sample a pair of ordinates, *K*_1_ *∼* U(*K*_min_, *K*_max_) and *K*_2_ *∼* U(*K*_min_, *K*_max_).
3. Repeat sampling pairs of ordinates until a pair of points on different sides of the threshold is identified, 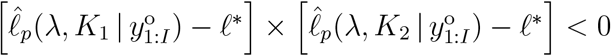.
4. Given these two points, use the bisection algorithm to estimate *K* such that 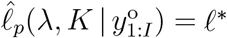.
5. Repeat the procedure of fixing *λ* and using the bisection algorithm to estimate *K* is *N* times.
6. To obtain a reasonable spread of points around the contour we repeat this procedure by first fixing *K* constant, and then using the bisection algorithm to estimate *λ* a further *N* times. This procedure identifies 2*N* discrete points on the 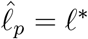 contour.

**Figure 4:**
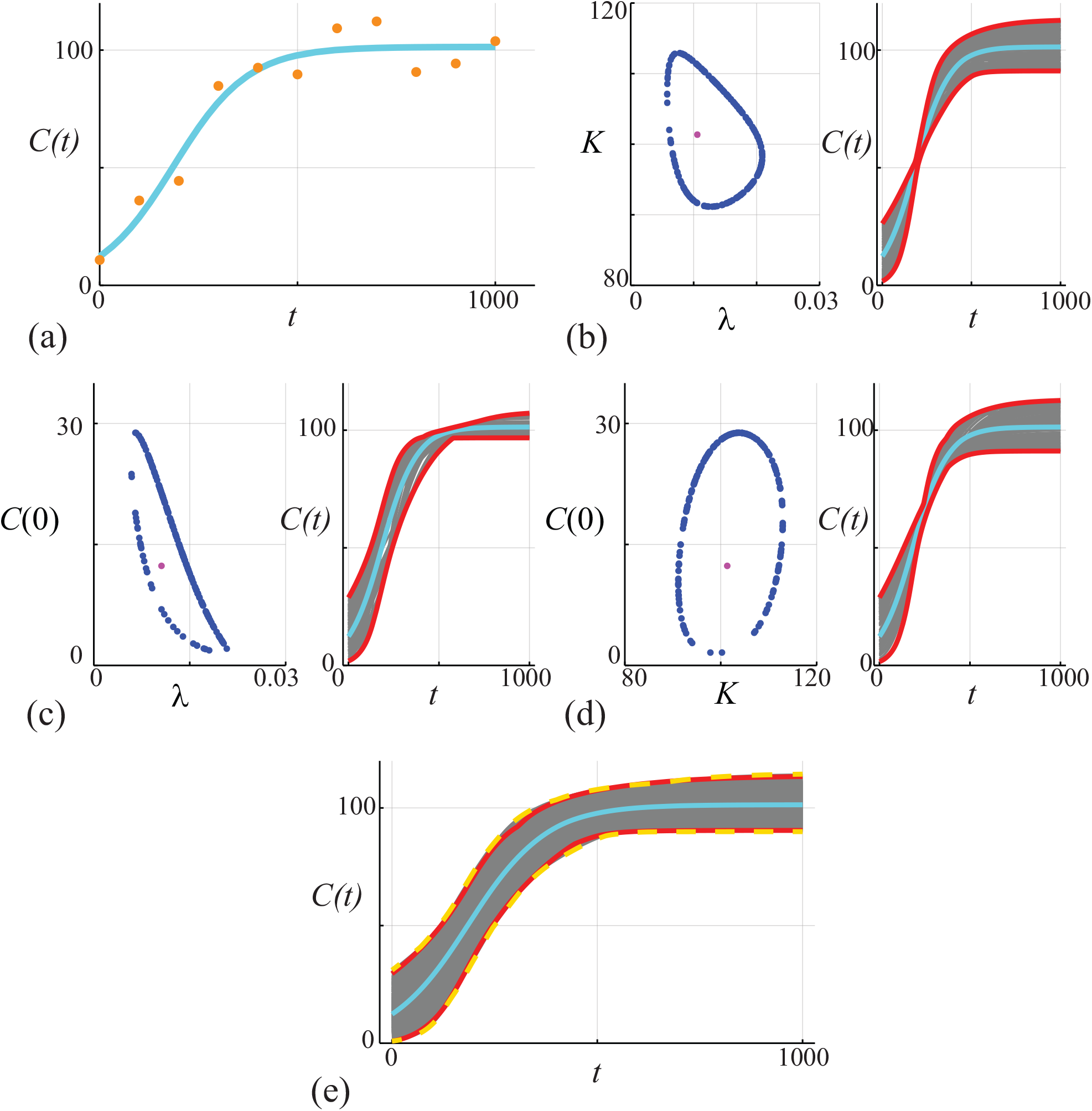
(a) Data obtained by solving Eq (22), with *θ* = (*λ, K, C*(0)) = (0.01, 100, 10), at *t* = 0, 100, 200, …, 1000 is corrupted with Gaussian noise with *σ* = 10. The MLE solution (cyan) is superimposed, 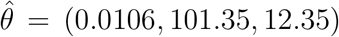. (b)-(d) Bivariate profiles for (*λ, K*), (*λ, C*(0)) and (*C*(0), *K*), respectively. In (b)-(d) the left-most panel shows 200 points along the 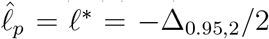 contours (blue discs) superimposed with the MLE (pink disc). The right-most panels shows 200 *C*(*t*) predictions (grey) that are associated with the 200 points along the 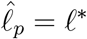 boundary, together with the profile-wise prediction interval (solid red). (e) Compares approximate prediction intervals obtained by computing the union of the profile-wise prediction intervals with the prediction intervals from the full likelihood function. Predictions (grey) are constructed by considering *N* = 10^4^ choices of *θ* and we plot solutions with 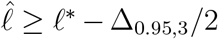 only. These solutions define a gold-standard prediction interval (dashed gold) that we compare with the MLE solution (cyan) and with the approximate prediction interval formed by the union of the three bivariate trajectories (red).

This procedure identifies 2*N* discrete points on the 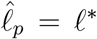 contour. While there are many other approaches to estimate and visualise the bivariate profile likelihood, we find this approach is both straightforward to implement and computationally robust.

Results in Figure 4(b)-(d) shows how our method identifies points along the boundary of the bivariate profile (blue discs), and for each boundary point identified we solve Eq (22) to give a family of pairwise predictions, as shown (grey solid curves). Taking the union of these pairwise profiles gives a prediction interval in Figure 4(e) (dashed red curves) that compares extremely well with the prediction interval estimated from the full likelihood function (yellow solid curves). Therefore, this boundary tracing method outperforms the results in Figures 2 and 3 and so we now focus on this approach for the remainder of this work. We caution, however, that this method assumes that the extreme points of the parameter confidence set determine the extreme points of the prediction (trajectory) confidence set.

### Case Study 2: Lotka-Volterra model of ecological competition with Gaussian noise

We now apply the same techniques to the ubiquitous Lotka-Volterra model of ecological competition [68], which describes the interactions between prey species *a*(*t*) with predator species *b*(*t*). This model can be written in non-dimensional form as

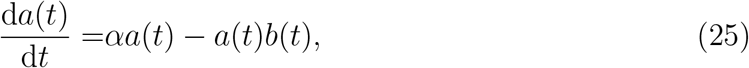

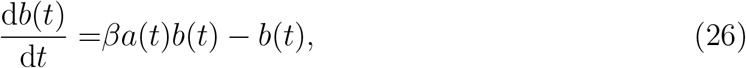

so that we have *θ* = (*α, β, a*(0), *b*(0)). In this example we assume that our observations consist of a time series of both species, i.e. 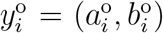, meaning that at each discrete time point we have an observation of both the prey and predator species. Further, we assume that, for all parameter sets, the observations of both the predator and prey are normally distributed about the solution *z*(*t*; *θ*) = (*a*(*t*; *θ*), *b*(*t*; *θ*)) of Eqs (25)–(26) with a common pre-specified constant variance, 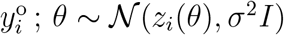. Here *I* is a 2 *×* 2 identity matrix and *z*_*i*_(*θ*) = *z*(*t*_*i*_; *θ*) = (*a*(*t*_*i*_; *θ*), *b*(*t*_*i*_; *θ*)). For this case study, we consider the solution of Eqs (25)–(26) on the interval 0 *≤ t ≤* 10, with 21 equally-spaced observations at *t*_*i*_ = (*i −* 1)*/*2 for *i* = 1, 2, 3, … 21. To make a distinction between estimation and prediction we use the first 15 time points (i.e. *t ≤* 7) for parameter estimation, whereas we make predictions over time whole time interval (i.e. *t ≤* 10).

Results in Figure 5(a) show 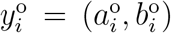 and the associated MLE solution, again noting that the total time series consists of 21 observations of each species, but here we only use the first 15 points in the time series for parameter estimation. In both Figure 5(a)-(b) we show a vertical dashed line at *t* = 7 to make it clear that data for *t ≤* 7 are used for model inference, whereas predictions are made over the whole time interval, *t ≤* 10.

**Figure 5:**
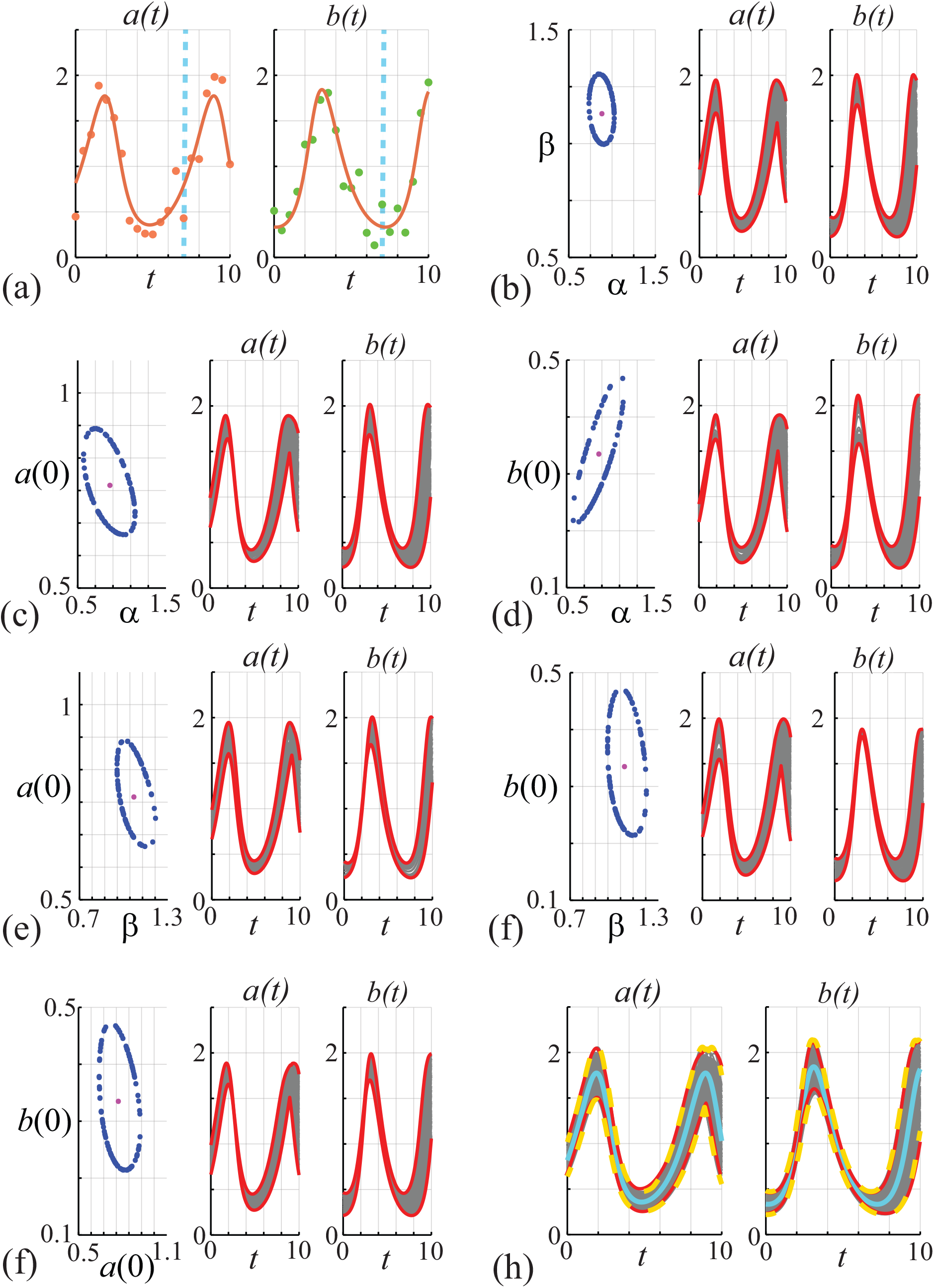
(a) Data obtained by solving Eqs (25)–(26), with *θ* = (*α, β, x*(0), *y*(0)) = (0.9, 1.1, 0.8, 0.3), at *t* = 0, 0.5, 1.0, …, 10 is corrupted with Gaussian noise with *σ* = 0.2. The MLE solution (solid curves) is superimposed, 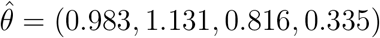. (b)-(g) Bivariate profiles for (*α, β*), (*α, x*(0)), (*α, y*(0)), (*β, x*(0)), (*β, y*(0)) and (*x*(0), *y*(0)), respectively. In (b)-(g) the left-most panel shows 100 points along the 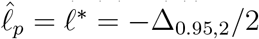 contour (blue discs) and the MLE (pink disc). The middle- and right-most panels show 100 predictions of *a*(*t*) and *b*(*t*), respectively, associated with the 100 points along the 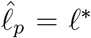 contour, together with the profile-wise prediction interval (solid red). (e) Compares approximate prediction intervals obtained by computing the union of the profile-wise prediction intervals with the prediction intervals from the full likelihood function.

Results in Figure 5(b)-(g) show a sequence of bivariate profiles and the associated profile-wise prediction intervals. The left-most panel in each of these plots show the MLE (pink discs) and 100 points along the 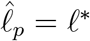 boundary identified using the method outlined for the results in Figure 4 (blue discs). The middle- and right-most panels in Figure 5(b)-(g) show solutions of Eqs (25)–(26) associated with the 100 points identified in the bivariate profiles along with the pairwise prediction interval (grey solid curves). As for the previous case study, the shape of the 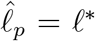 boundary identifies potential relationships between the pairs of parameters in terms of their effects on model trajectories, and the profile-wise prediction intervals for each parameter pair provide insight in the how uncertainty in these parameter estimates translates into uncertainty in the solution of the mathematical models (red solid curves). Results in Figure 5(h) compares the prediction interval formed by taking the union of the profile-wise prediction sets (dashed red curves) with the prediction interval taken from working with the full likelihood function (solid yellow curves). As with the simpler logistic model, we see that the union of pairwise profiles provides a relatively accurate alternative to working with the full likelihood function.

### Case Study 3: Logistic growth with Poisson noise

Here we return to the dimensional logistic growth model, Eq (22) with *θ* = (*λ, K, C*(0)), except that we work with a non-normal Poisson noise model by assuming that

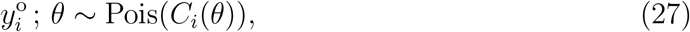

which is often used to model non-negative quantities, such as population densities in ecological modelling [76–79]. Despite the difference in the observation model, the approximate calibration criterion for the likelihood function for the Poisson noise model is the same as before [27]. The results in Figure 6(a) show data and the MLE solution of Eq (22). Comparing the data in Figure 2(a) and Figure 6(a) we see that with the Poisson noise model we have larger variability as *C*(*t*) increases, whereas with the Gaussian noise model we have approximately constant variability over time. Working with the non-Gaussian noise model means that the variability vanishes as *C*(*t*) *→* 0^+^ which avoids the possibility of negative population densities. Results in Figure 6(b) show the associated univariate profiles (solid blue curves) where we see that each profile is well-formed about a single peak. Using the same procedure as in Figure 2, we generate the profile-wise prediction sets shown in Figure 6(c)-(e), where we see that we have the same qualitative trends in terms of how variability in each parameter impacts the prediction interval for *C*(*t*). Taking the union of the profile-wise prediction sets leads to the prediction set in Figure 6(f) (dashed red curves), which, when compared to the prediction interval obtained using the full likelihood (solid yellow curves), leads to an approximation of a gold-standard prediction interval that underestimates the uncertainty and motivates us again to consider working with the pairwise bivariate profiles.

**Figure 6:**
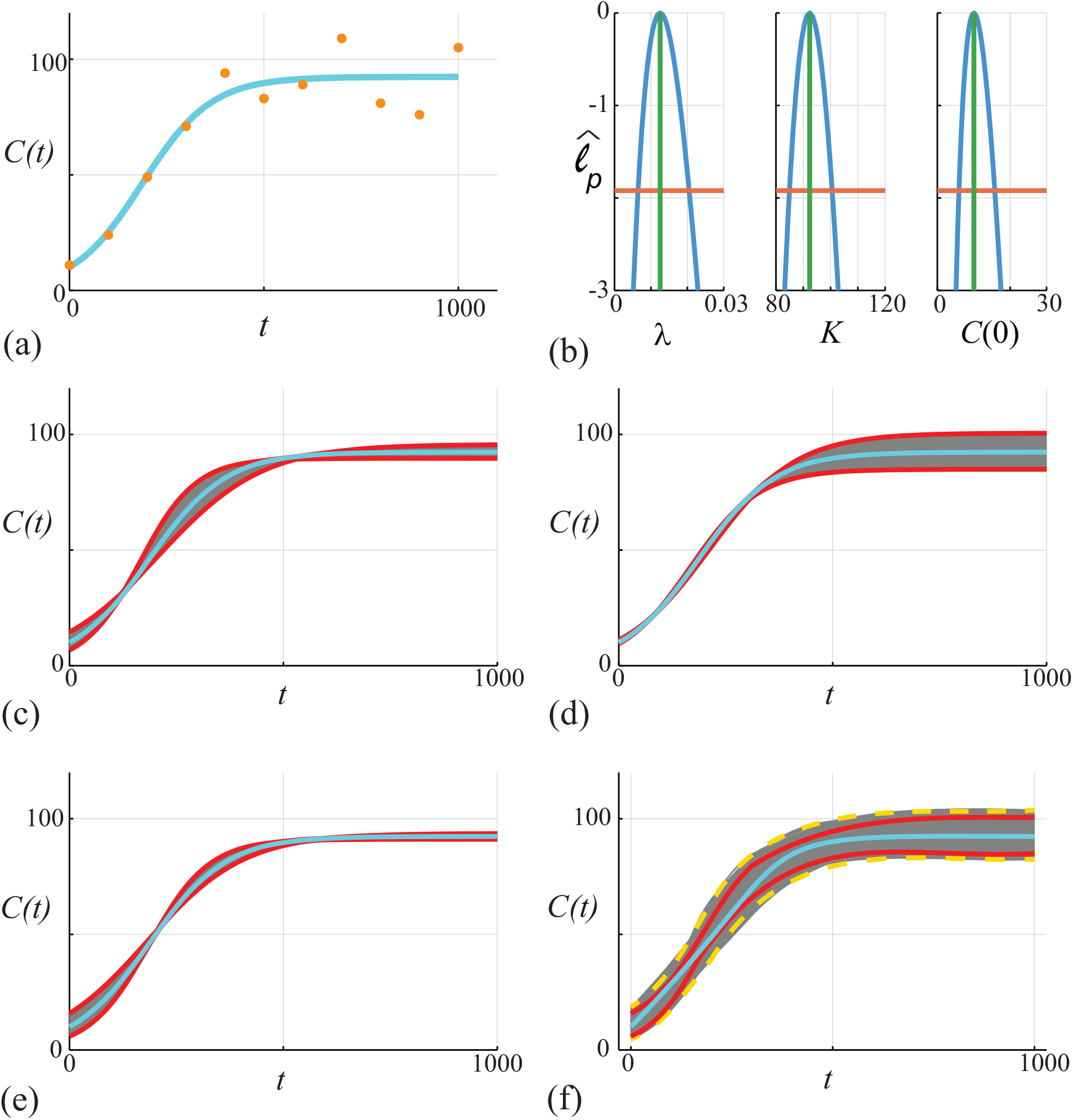
(a) Data obtained by solving Eq (22), with *θ* = (*λ, K, C*(0)) = (0.01, 100, 10), at *t* = 0, 100, 200, …, 1000 is corrupted with Poisson noise. The MLE solution (cyan) is superimposed, 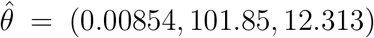. (b) Univariate profiles for *λ, K* and *C*(0), respectively. Each profile is superimposed with the MLE (vertical green) and the 95% threshold 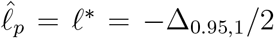 is shown (horizontal orange). Points of intersection of each profile and the threshold define asymptotic confidence intervals: 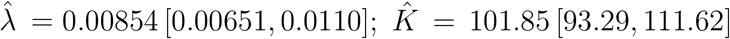 and, 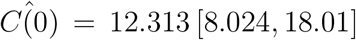. (c)-(e) *C*(*t*) predictions associated with the *λ, K* and *C*(0) univariate profiles in (b), respectively. In each case we uniformly discretise the interest parameter using *N* = 100 points along the 95% confidence interval identified in (b), and solve the model forward with *θ* = (*ψ, ω*) and plot the family of solutions (grey), superimposing the MLE (cyan), and we identify the prediction interval defined by these solutions (solid red). (f) Compares approximate prediction intervals obtained by computing the union of the three univariate profiles with the prediction intervals computed from the full likelihood. Trajectories (grey) are constructed by considering *N* = 10^4^ choices of *θ* and we plot solutions with *ℓ ≥ ℓ*^*∗*^ = − ∆_0.95,3_*/*2 only. These solutions define a prediction interval (dashed gold) that we compare with the MLE solution (cyan) and with the approximate prediction interval formed by taking the union of the three univariate intervals (red).

Results in Figure 7 are directly analogous to those in Figure 3 except that here we work with the Poisson noise model. Results in the left-most panels in Figure 7(b)-(d) show 200 points on the 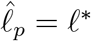 boundaries identified as in the previous case studies (blue discs). As we saw with the Gaussian noise model previously, the shape of the identified boundaries identifies relationships between pairs of parameters in terms of their effects on the model solution. The right-most panels in Figure 7(b)-(d) shows the solution of Eq (22) corresponding to the points along the *ℓ* = *ℓ*^*∗*^ boundaries (grey solid curves). As for the results with the Gaussian noise model in Figure 3, the union of the profile-wise predictions in Figure 7(e) (dashed red curves) compares very well with the prediction interval obtained using the full likelihood function (yellow solid curves).

**Figure 7:**
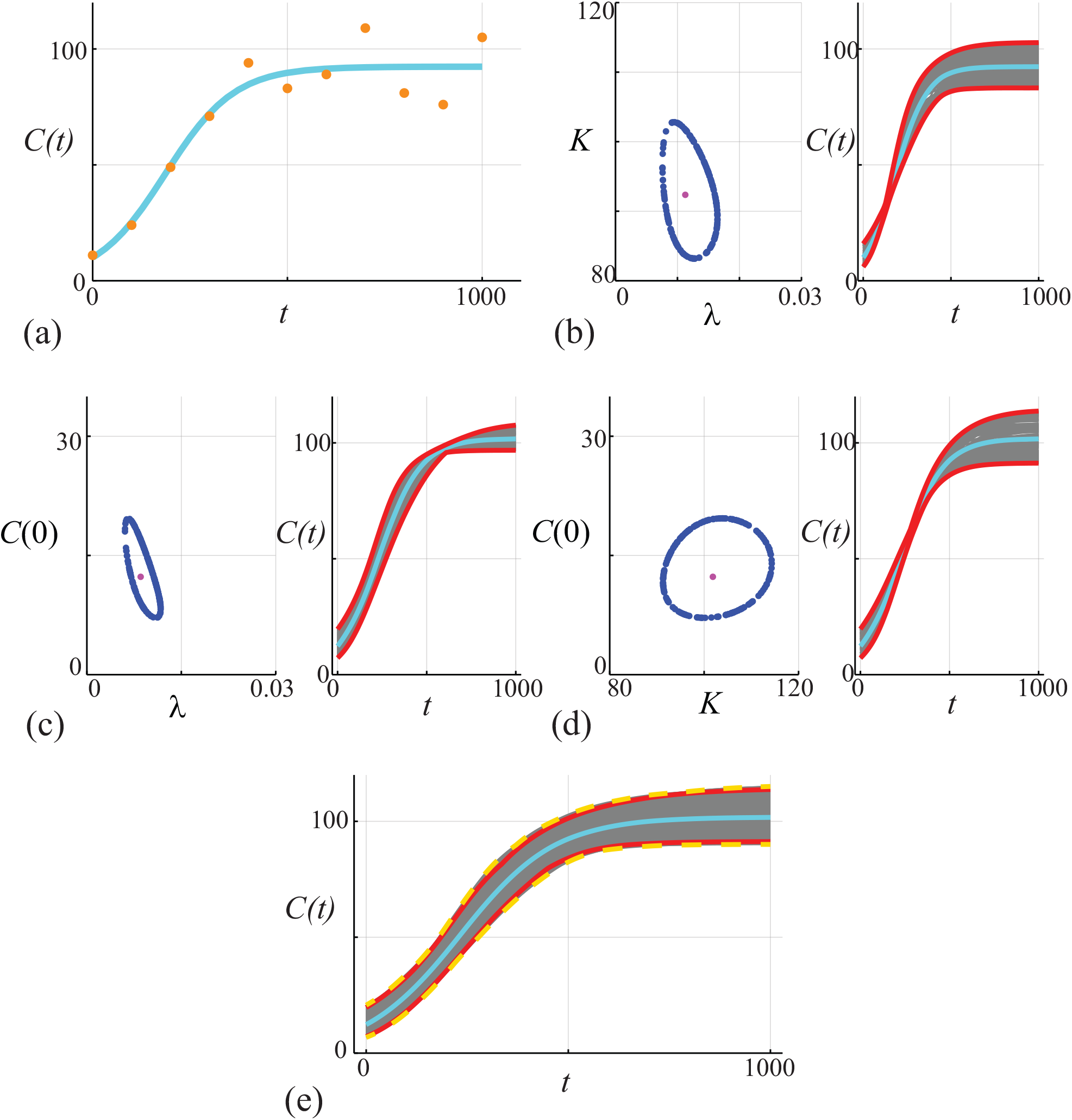
(a) Data obtained by solving Eq (22), with *θ* = (*λ, K, C*(0)) = (0.01, 100, 10), at *t* = 0, 100, 200, …, 1000 is corrupted with Poisson noise. The MLE solution (cyan) is superimposed, 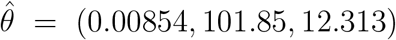. (b)-(d) Bivariate profiles for (*λ, K*), (*λ, C*(0)) and (*C*(0), *K*), respectively. The left-most panel in (b)-(d) shows 200 points along the 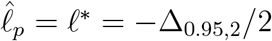 contour (blue discs) and the MLE (pink disc). The rightmost panel in (b)-(d) shows 200 *C*(*t*) predictions associated with the points along the 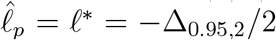 contour that define a prediction interval (solid red). (e) Compares approximate prediction intervals obtained by computing the union of the three bivariate profiles with the exact prediction interval computed from the full likelihood. Trajectories (grey) are constructed by considering *N* = 10^4^ choices of *θ* and we plot solutions with 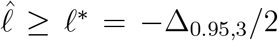 only. These solutions define an exact prediction interval (dashed gold) that we compare with the MLE solution (cyan) and with the approximate prediction interval formed by taking the union of the three univariate intervals (red).

While this case study focuses on a Poisson noise model in the context of ecological population dynamics [76–79], other kinds of non-Gaussian noise models can also be accommodated within the workflow [69]

## Discussion

This work outlines a likelihood-based workflow, called Profile-Wise Analysis, for identifiability analysis, estimation, and prediction for mechanistic mathematical models. Three case studies are presented to make our theoretical developments as practical as possible: a single nonlinear ODE mechanistic model with a Gaussian noise model; (ii) a system of coupled nonlinear ODEs with a Gaussian noise model; and (iii) a single nonlinear ODE mechanistic model with non-Gaussian noise.

While our workflow combines parameter identifiability, parameter estimation, and model prediction, we intentionally focus much of the theoretical development and our case studies on linking parameter estimates to data distribution parameters representing model predictions. We choose this emphasis as prediction questions often receive less attention in the likelihood-based setting than questions about parameter estimation and identifiability. Furthermore, linking parameter estimation back to prediction naturally completes the analysis cycle and provides the opportunity for iterative model building, testing and refining. Our primary predictive quantity of interest here is the model trajectory, intended to represent the data mean. For completeness, we also sketch a method for predicting data realisations that leverages our ability to form and propagate confidence sets for both mechanistic model and data distribution parameters. However, the proper evaluation and comparison of such methods for predicting realisations rather than model trajectories remain for future work.

A central component of our workflow is the use of profile likelihood to construct profile-wise prediction intervals that propagate confidence sets for model parameters forward to data distribution parameters in a way that explicitly isolates how different parameter combinations affect predictions [23, 24]. We demonstrate this important step using various types of approximation. First, we consider univariate profile likelihood functions and explicitly propagate parameter sets forward to show how individual parameters affect model trajectories [23, 24]. Using individual profiles is insightful because it allows us to explore how particular parameters, describing different mechanisms inherent in the mathematical models, affects various aspects of the model predictions. Taking the union of the single-parameter profile-wise prediction intervals also allows us to construct an approximate overall prediction interval for the model trajectory that incorporates the effects of each parameter. However, we find that such intervals are typically narrower than the prediction intervals obtained by working with the full likelihood, leading to potential under-coverage and underestimation of uncertainties. This leads us to consider working with bivariate profiles rather than univariate profiles, both for estimation and for taking the union of bivariate profile-wise prediction intervals. By working with bivariate profile likelihood functions, we explicitly see how pairs of parameters jointly affect predictions, offering insight into synergistic effects. Furthermore, taking the union of these two-dimensional profile-wise prediction intervals leads to combined prediction intervals that compare very well with results obtained by working with the full likelihood function. We consider both grid-based and boundary-based methods of computation, finding that the latter does a better job of capturing non-elliptical profile likelihood functions.

Working within an ‘ODE + noise’ framework here is natural for the purposes of presenting and demonstrating the workflow since this is a very common situation in practice. However, our framework can be applied to other situations. More broadly, the workflow can also apply to other differential equation-based mechanistic models, such as partial differential equation-based models [13], stochastic differential equation-based models [63], as well as simulation-based models [34, 35, 41, 80, 81]. Regardless of the form of the mechanistic model, the critical requirement of our framework is that we have access to a likelihood function. In the case of a simulation-based stochastic model, we assume an auxiliary likelihood function [31], also known as a synthetic or surrogate likelihood [32, 33], is available. For example, our recent work [35] considered a random walk simulation-based model of diffusive heat transfer across layered heterogeneous materials, and we generated data in terms of breakthrough-type time series to mimic experiments involving heat transfer across layered skin. For identifiability analysis and inference, we made use of an approximate likelihood function based on moment matching, written in terms of a Gamma distribution, and showed that this led to accurate inference at a fraction of the computational cost of working directly with the stochastic random walk simulations. Similarly, we recently demonstrated how to approximate stochastic individual-based models using models of the form ‘ODE + noise’ by appropriately correcting the noise model using surrogate modelling through coarse graining [34]. Therefore, simulation-based mechanistic model approaches fall naturally within our workflow when a synthetic or surrogate like-lihood function can be constructed. However, additional investigation into the accuracy of such approximations is needed, e.g., as in other work on Monte Carlo profile likelihood methods [41]. In this work we deliberately focused the first two case studies on situations involving pre-specified Gaussian noise with constant variance since this is very common in practice [13, 70]. Other noise models can be accommodated within the workflow such as working with a Gaussian noise model with unknown constant variance [24], a prespecified Gaussian noise model with variable variance [82], or non-Gaussian noise models as we illustrate with the third case study.

## Conclusions and future directions

An important feature of our work here is that all three case studies involve practically identifiable models, and we have yet to directly address the question of applying our complete workflow to problems that are structurally unidentifiable [44, 83, 84]. Common motivations for assessing structural identifiability of mechanistic models are: to use this as a screening tool before more expensive estimation is attempted [13, 48], and to simplify structurally unidentifiable models to structurally identifiable models to obtain more mechanistic insight [85, 86]. While profile likelihood, as used in our present workflow, is established as a valuable tool for initial parameter identifiability analysis capable of detecting non-identifiable models [44, 48, 55], we have not explicitly considered the consequences of structural non-identifiability for predictions and model simplification. However, we are also interested in such scenarios, and we leave these for future consideration where it is likely that we would need to use a different numerical optimization approach, such as using global search algorithms instead of local strategies. Another point of interest for future exploration is to examine different options for constructing the profile-wise predictions. In this work we take a very straightforward approach of using a uniform mesh and identifying the boundary using a simple bisection algorithm. Other options for sampling the parameter space (e.g. random samples or Latin Hypercube-based samples) or tracing the boundary (e.g. gradient-based methods using automatic differentiation and continuation methods) are available, and a detailed comparison of different methodologies would be insightful. In this work we have been careful to compare prediction intervals for our approximate method with prediction intervals obtained by working with the full-likelihood, as the full likelihood is a gold standard likelihood-based approach. It would also be useful to directly compare the new approximate method with other established approximate frequentist methods, e.g. as reviewed in [15], as well as working with established benchmark problems from the systems biology literature, e.g. as reviewed in [87]. However, in this first presentation of our PWA approach we have restricted our attention to relatively simple, yet widely understood Case study problems of broad interest across the mathematical biology and mathematical ecology literature to emphasise the general nature of our approach.

## Acknowledgments

We thank Corey Yanofsky, Dominic Edelmann, and Thomas Lumley for useful conversations about confidence intervals and simultaneous inference.

## References

[1] Chatzilena A, van Leeuwen E, Ratmann O, Baguelin M, Demiris N. (2019). Contemporary statistical inference for infectious disease models using Stan. Epidemics, 29, 100367. (doi: 10.1016/j.epidem.2019.100367).

[2] Gabry J, Simpson D, Vehtari A, Betancourt M, Gelman A. 2019. Visualization in Bayesian workflow. Journal of the Royal Statistical Society: Series A (Statistics in Society), 182(32). (doi: 10.1111/rssa.12378).

[3] Gelman A. 2004. Exploratory data analysis for complex models. Journal of Computational and Graphical Statistics. 13, 755–779. (doi: 10.1198/106186004X11435).

[4] Gelman A, Carlin JB, Stern HS, Dunson DB, Vehtari A, Rubin DB. 2013. Bayesian data analysis, third edition. Taylor & Francis.

[5] Gelman A, Vehtari A, Simpson D, Margossian CC, Carpenter B, Yao Y, Kennedy L, Gabry J, Bürkner PC, Modrák M. 2020. Bayesian workflow. arXiv preprint. (arxiv.org/abs/2011.01808).

[6] Grinsztajn L, Semenova E, Margossian CC, Riou J. 2021. Bayesian workflow for disease transmission modeling in Stan. Statistics in medicine, 40(27), 6209–6234. (doi: 10.1002/sim.9164).

[7] Sunnåker M, Busetto AG, Numminen E, Corander J, Foll M, Dessimoz C. 2013. Approximate Bayesian Computation. PLOS Computational Biology. 9: e1002803. (doi: 10.1371/journal.pcbi.1002803).

[8] Wasserman L. 2004. All of statistics: a concise course in statistical inference. Springer.

[9] Cox DR. 2006. Principles of statistical inference. Cambridge University Press.

[10] Neyman J 1977. Frequentist probability and frequentist statistics. Synthese, 97–131.

[11] Bernardo JM, Smith AF. 2009. Bayesian theory (Vol. 405). John Wiley & Sons.

[12] MacKay DJ 2003. Information theory, inference and learning algorithms. Cambridge university press.

[13] Simpson MJ, Baker RE, Vittadello ST, Maclaren OJ. 2020. Parameter identifiability analysis for spatiotemporal models of cell invasion. Journal of the Royal Society Interface. 17, 20200055. (doi: 10.1098/rsif.2020.0055).

[14] Hinkley, D. 1979. Predictive likelihood. The Annals of Statistics, 7(4), 718–728.

[15] Villaverde AF, Raimúndez E, Hasenauer J, Banga JR. 2022. Assessment of prediction uncertainty quantification methods in systems biology. IEEE/ACM Transactions on Computational Biology and Bioinformatics. Early view. (doi:10.1109/TCBB.2022.3213914).

[16] Gutenkunst RN, Casey FP, Waterfall JJ, Myers CR, Sethna JP. (2007). Extracting falsifiable predictions from sloppy models. Annals of the New York Academy of Sciences, 1115(1), 203–211.

[17] Hass H, Kreutz C, Timmer J, Kaschek D. 2016. Fast integration-based prediction bands for ordinary differential equation models. Bioinformatics. 32, 1204–1210. (doi:10.1093/bioinformatics/btv743).

[18] Kreutz C, Raue A, Kaschek D, Timmer J. 2013. Profile likelihood in systems biology. The FEBS Journal. 280, 2564–2571. (doi:10.1111/febs.12276).

[19] Kreutz C, Raue A, Timmer J. 2013. Likelihood based observability analysis and confidence intervals for predictions of dynamics models. BMC Systems Biology. 6, 120. (doi:10.1186/1752-0509-6-120).

[20] Villaverde AF, Bongard S, Mauch K, Müller D, Balsa-Canto E, chmid J, Banga JR. 2015. A consensus approach for estimating the predictive accuracy of dynamic models in biology. Computer Methods and Programs in Biomedicine. 119, 17–28. (doi:10.1016/j.cmpb.2015.02.001).

[21] Oliver D, He N, Reynolds AC (1996) Conditioning permeability fields to pressure data. In Proc. 5th Eur. Conf. Mathematics of Oil Recovery, Sept.

[22] Wu D, Petousis-Harris H, Paynter J, Suresh V, Maclaren OJ. 2023. Likelihood-based estimation and prediction for a measles outbreak in Samoa. Infectious Disease Modelling. 8, 212–227. (doi: 10.1016/j.idm.2023.01.007).

[23] Murphy RJ, Maclaren OJ, Calabrese AR, Thomas PB, Warne DJ, Williams ED, Simpson MJ. 2022. Computationally efficient framework for diagnosing, understanding, and predicting biphasic population growth. Journal of the Royal Society Interface. 19, 20220560. doi: 10.1098/rsif.2022.0560.

[24] Simpson MJ, Walker SA, Studerus EN, McCue SW, Murphy RJ, Maclaren OJ. 2022. Profile likelihood-based parameter and predictive interval analysis guides model choice for ecological population dynamics. Mathematical Biosciences. 355, 108950. 10.1016/j.mbs.2022.108950.

[25] Maclaren OJ, Nicholson R. 2019. What can be estimated? Identifiability, estimability, causal inference and ill-posed inverse problems. arXiv. https://arxiv.org/abs/1904.02826.

[26] Maclaren, OJ, Nicholson, R. 2021. Models, identifiability, and estimability in causal inference. In 38th International Conference on Machine Learning. Workshop on the Neglected Assumptions in Causal Inference. ICML.

[27] Pawitan Y. 2001. In all likelihood: statistical modelling and inference using likelihood. Oxford University Press.

[28] Sprott DA. 2008. Statistical inference in science. Springer Science & Business Media.

[29] Cranmer K, Brehmer J, Louppe G. 2020. The frontier of simulation-based inference. Proceedings of the National Academy of Sciences, 117(48), 30055–30062. (doi: 10.1073/pnas.1912789117).

[30] Diggle, PJ., Gratton, R. J. 1984. Monte Carlo methods of inference for implicit statistical models. Journal of the Royal Statistical Society: Series B (Methodological), 46(2), 193–212.

[31] Drovandi CC, Pettitt AN, Lee A. 2015. Bayesian indirect inference using a parametric auxiliary model. Statistical Science, 30(1), 72–95.

[32] Fasiolo M, Pya N, Wood SN. 2016. A comparison of inferential methods for highly nonlinear state space models in ecology and epidemiology. Statistical Science, 96–118. (http://www.jstor.org/stable/24780835).

[33] Wood SN. 2010. Statistical inference for noisy nonlinear ecological dynamic systems. Nature. 466, 1102–1104. (doi: 10.1038/nature09319).

[34] Simpson MJ, Baker RE, Buenzli PR, Nicholson R, Maclaren, OJ. 2022. Reliable and efficient parameter estimation using approximate continuum limit descriptions of stochastic models. Journal of Theoretical Biology. 549, 111201. (doi: 10.1016/j.jtbi.2022.111201).

[35] Simpson MJ, Browning AP, Drovandi C, Carr EJ, Maclaren OJ, Baker RE. 2021. Profile likelihood analysis for a stochastic model of diffusion in heterogeneous media. Proceedings of the Royal Society A: Mathematical, Physical and Engineering Sciences. 477, 20210214. (doi:10.1098/rspa.2021.0214).

[36] Heggland K, Frigessi A. 2004. Estimating functions in indirect inference. Journal of the Royal Statistical Society: Series B (Statistical Methodology), 66(2), 447–462. (http://www.jstor.org/stable/3647536).

[37] Gourieroux, C, Monfort, A, Renault, E. 1993. Indirect inference. Journal of Applied Econometrics. 8, S85–S118. (http://www.jstor.org/stable/2285076).

[38] Simpson MJ, Browning AP, Warne DJ, Maclaren OJ, Baker RE. 2022. Parameter identifiability and model selection for sigmoid population growth models. Journal of Theoretical Biology. 535, 110998. (doi: 10.1016/j.jtbi.2021.110998).

[39] Ong VMH, Nott DJ, Tran MN, Sisson SA, Drovandi CC. 2018. Likelihood-free inference in high dimensions with synthetic likelihood. Computational Statistics & Data Analysis, 128. 271–291. doi: 10.1016/j.csda.2018.07.008.

[40] Price LF, Drovandi CC, Lee A, Nott DJ (2018). Bayesian synthetic likelihood. Journal of Computational and Graphical Statistics. 27, 1–11. doi: 10.1080/10618600.2017.1302882.

[41] Ionides, EL, Breto, C, Park, J, Smith, RA, King, AA. 2017. Monte Carlo profile confidence intervals for dynamic systems. Journal of The Royal Society Interface, 14(132), 20170126. (doi: 10.1098/rsif.2017.0126).

[42] Pace L, Salvan A. 1997. Principles of Statistical Inference from a Neo-Fisherian Perspective. In: Advanced Series on Statistical Science and Applied Probability, vol. 4. World Scientific, Singapore.

[43] Campbell DA, Chkrebtii O. 2013. Maximum profile likelihood estimation of differential equation parameters through model-based smoothing state estimate. Mathematical Biosciences. 246, 283–292. (doi:10.1016/j.mbs.2013.03.011).

[44] Chiş O, Villaverde AF, Banga JR, Balsa-Canto E. 2016. On the relationship between sloppiness and identifiability. Mathematical Biosciences. 282, 147–161. (doi: 10.1016/j.mbs.2016.10.009).

[45] Eisenberg MC, Hayashi MAL. 2014. Determining identifiable parameter combinations using subset profiling. Mathematical Biosciences. 256, 115–126. (doi:10.1016/j.mbs.2014.08.008).

[46] Cole D. 2020. Parameter redundancy and identifiability. CRC Press.

[47] Raue A, Kreutz C, Maiwald T, Bachmann J, Schilling M, Klingmüller U, Timmer J. 2009. Structural and practical identifiability analysis of partially observed dynamical models by exploiting the profile likelihood. Bioinformatics. 25, 1923–1929. (doi: 10.1093/bioinformatics/btp358).

[48] Raue A, Kreutz C, Theis FJ, Timmer J. 2013. Joining forces of Bayesian and frequentist methodology: a study for inference in the presence of non-identifiability. Philosophical Transactions of the Royal Society A: Mathematical, Physical and Engineering Sciences. 371, 20110544. (doi: 10.1098/rsta.2011.0544).

[49] Raue A, Karlsson J, Saccomani MP, Jirstrand M, Timmer J. 2014. Comparison of approaches for parameter identifiability analysis of biological systems. Bioinformatics. 30, 1440–1448. (doi: 10.1093/bioinformatics/btu006).

[50] Murphy RJ, Gunasingh G, Haass NK, Simpson MJ. 2023. Growth and adaptation mechanisms of tumour spheroids with time-dependent oxygen availability. PLOS Computational Biology. 19, e1010833. (doi: 10.1371/journal.pcbi.1010833).

[51] Wieland F-G, Hauber AL, Rosenblatt M, Tönsing C, Timmer J. 2021. On structural and practical identifiability. Current Opinion in Systems Biology. 25, 60–69. (doi:10.1016/j.coisb.2021.03.00).

[52] Browning AP, Drovandi C, Turner IW, Jenner AL, Simpson MJ (2022) Efficient inference and identifiability analysis for differential equation models with random parameters. PLOS Computational Biology. 18: e1010734. (doi: 10.1371/journal.pcbi.1010734).

[53] Siekmann I, Sneyd J, Crampin EJ. 2012. MCMC can detect nonidentifiable models. Biophysical Journal. 103, 2275–2286. (doi: 10.1016/j.bpj.2012.10.024).

[54] Siekmann I, Fackrell M, Crampin EJ, Taylor P. 2016. Modelling modal gating of ion channels with hierarchical Markov models. Proceedings of the Royal Society A: Mathematical, Physical and Engineering Sciences. 472, 20160122. (doi:10.1098/rspa.2016.0122).

[55] Fröhlich F, Theis FJ, Hasenauer J. 2014. Uncertainty analysis for non-identifiable dynamical systems: Profile likelihoods, bootstrapping and more. International Conference on Computational Methods in Systems Biology. 61–72. Springer.

[56] Edwards, AWF. 1969. Statistical methods in scientific inference, Nature 222, 1233–1237.

[57] Theil H, Goldberger AS. 1961. On pure and mixed statistical estimation in economics. International Economic Review. 2, 65–78. doi: 10.2307/2525589.

[58] Stark PB. 2015. Constraints versus priors. SIAM/ASA Journal on Uncertainty Quantification. 3, 586–598. (doi: 10.1137/130920721).

[59] Patel JK. 1989. Prediction intervals-a review. Communications in Statistics-Theory and Methods. 18, 2393–2465. doi: 10.1080/03610928908830043.

[60] Tian Q, Nordman DJ, Meeker WQ. 2022. Methods to compute prediction intervals: A review and new results. Statistical Science. 37, 580–597. doi: 10.1214/21-STS842.

[61] Sulieman H, McLellan PJ, Bacon DW. 2001. A profile-based approach to parametric sensitivity analysis of nonlinear regression models. Technometrics. 43, 425–433. http://www.jstor.org/stable/1270813.

[62] Sulieman, H, McLellan, PJ, Bacon, DW. 2004. A profile-based approach to parametric sensitivity in multiresponse regression models. Computational Statistics & Data Analysis, 45(4), 721–740. doi: 10.1198/00401700152672519.

[63] Browning AP, Warne DJ, Burrage K, Baker RE, Simpson MJ (2020). Identifiability analysis for stochastic differential equations models in systems biology. Journal of the Royal Society Interface. 17, 20200652. (doi:10.1098/rsif.2020.0652).

[64] Juul JL, Græsbøll K, Christiansen LE, Lehmann S. 2021. Fixed-time descriptive statistics underestimate extremes of epidemic curve ensembles. Nature Physics. 17, 5–8. (doi: 10.1038/s41567-020-01121-y).

[65] Miller RGJ. 1981. Simultaneous Statistical Inference. 2nd edn, Springer.

[66] Lieberman GJ, Miller RG. 1963. Simultaneous tolerance intervals in regression. Biometrika. 50, 155–168. (doi:10.2307/2333756).

[67] Casella G, Berger R. 2001. Statistical Inference. Belmont, CA: Duxbury.

[68] Toni T, Welch D, Strelkowa N, Ipsen A, Stumpf MPH. 2009. Approximate Bayesian computation scheme for parameter inference and model selection in dynamical systems. Journal of the Royal Society Interface. 6, 187–202. (doi:10.1098/rsif.2008.0172).

[69] He D, Ionides EL, King AA. 2010. Plug-and-play inference for disease dynamics: measles in large and small populations as a case study. Journal of the Royal Society Interface. 7, 271–283. (doi: 10.1098/rsif.2009.0151).

[70] Hines KE, Middendorf TR, Aldrich RW. 2014. Determination of parameter identifiability in nonlinear biophysical models: A Bayesian approach. Journal of General Physiology. 143, 401. (doi: 10.1085/jgp.201311116).

[71] Tsoularis A, Wallace J. 2002. Analysis of logistic growth models. Mathematical Biosciences. 179, 21–55. (doi:10.1016/S0025-5564(02)00096-2).

[72] Steele J, Adams J, Slukin T. 1998 Modelling paleoindian dispersals. World Archaeology. 30, 286-305. 10.1080/00438243.1998.9980411.

[73] Murphy RJ, Maclaren OJ, Simpson MJ. 2023. Implementing measurement error models for estimation and prediction in the life sciences. arXiv preprint. (doi:10.48550/arXiv.2307.01539).

[74] Rackauckas C, Nie Q. 2017. DifferentialEquations.jl - a performant and feature-rich ecosystem for solving differential equations in Julia. Journal of Open Research Software. 5, 15. (doi: 10.5334/jors.151).

[75] Johnson SG. 2022. The NLopt module for Julia. Retrieved May 2023 NLopt.

[76] Auger-Méthé M, Newman K, Cole D, Empacher F, Gryba R, King AA, Leos-Barajas V, Flemming JM, Nielsen A, Petris G, Thomas L. 2021. A guide to state–space modeling of ecological time series. Ecological Monographs. 91, e01470. (doi:10.1002/ecm.1470).

[77] Hefley TJ, Tyre AJ, Blankenship EE. 2013. Statistical indicators and state–space population models predict extinction in a population of bobwhite quail. Theoretical Ecology. 6, 319. (doi: 10.1007/s12080-013-0195-3).

[78] Hefley TJ, Tyre AJ, Blankenship EE. 2013. Fitting population growth models in the presence of measurement and detection error. Ecological Modelling. 263, 244–250. (doi: 10.1016/j.ecolmodel.2013.05.003).

[79] Knape J, Jonzen N, Skold M. 2011. On observation distributions for state space models of population survey data. Journal of Animal Ecology 2011. 80, 1269–1277. (doi: 10.1111/j.1365-2656.2011.01868.x).

[80] Breto C. 2018. Modeling and inference for infectious disease dynamics: a likelihood-based approach. Statistical Science. 33, 57–69. (doi:10.1214/17-STS636).

[81] Breto C, Ionides EI, King AA. 2019. Panel data analysis via mechanistic models. Journal of the American Statistical Association. 115, 1178–1188. (doi:10.1080/01621459.2019.1604367).

[82] Browning AP, Maclaren OJ, Lanaro M, Allenby MC, Woodruff MA, Simpson MJ (2021). Model-based data analysis of tissue growth in thin 3D printed scaffolds. Journal of Theoretical Biology. 528: 110852. (doi:10.1016/j.jtbi.2021.110852).

[83] Chiş O, Banga JR, Balsa-Canto E. 2011. Structural identifiability of systems biology models: a critical comparison of methods. PLoS One. 6, e27755. (doi:10.1371/journal.pone.0027755).

[84] Chiş O, Banga JR, Balsa-Canto E. 2011. GenSSI: a software toolbox for structural identifiabilty analysis of biological models. Bioinformatics. 18, 2610–2611. (doi:10.1093/bioinformatics/btr431).

[85] Meshkat N, Eisenberg M, DiStefano III JJ. 2009. An algorithm for finding globally identifiable parameter combinations of nonlinear ODE models using Gröbner Bases. Mathematical Biosciences. 222, 61–72. (doi: 10.1016/j.mbs.2009.08.010).

[86] Meshkat N, Sullivan S, Eisenberg M. 2018. Identifiability results for several classes of linear compartment models. Bulletin of Mathematical Biology. 8, 1620–1651. (doi: 10.1007/s11538-015-0098-0).

[87] Hass H, Loos C, Raimúndez-Álvarez E, Timmer J, Hasenauer J, Kreutz C. 2019. Benchmark problems for dynamic modeling of intracellular processes. Bioinformatics. 35, 3073–3082. (doi:10.1093/bioinformatics/btz020).

